# Proteomic characterization of the Alzheimer’s disease risk factor BIN1 interactome

**DOI:** 10.1101/2025.03.09.642169

**Authors:** Joseph D. McMillan, Shuai Wang, Jessica Wohlfahrt, Jennifer Guergues, Stanley M. Stevens, Gopal Thinakaran

**Author notes:** Corresponding Author: **Gopal Thinakaran:**.

## Abstract

The gene *BIN1* is the second-largest genetic risk factor for late-onset Alzheimer’s disease (LOAD). It is expressed in neurons and glia in the brain as cell-type specific and ubiquitous isoforms. BIN1 is an adaptor protein that regulates membrane dynamics in many cell types. Previously, we reported that BIN1 predominantly localizes to presynaptic terminals in neurons and regulates presynaptic vesicular release. However, the function of neuronal BIN1 in relation to LOAD is not yet fully understood. A significant gap in the field is the unbiased characterization of neuronal BIN1-interacting proteins and proximal neighbors. To address this gap and help define the functions of neuronal BIN1 in the brain, we employed TurboID-based proximity labeling to identify proteins biotinylated by the neuronal BIN1 isoform 1-TurboID fusion protein (BIN1iso1-TID) in cultured mouse neuroblastoma (N2a) cells *in vitro* and in adult mouse brain neurons *in vivo*. Label-free quantification-based proteomic analysis of the BIN1iso1-TID biotinylated proteins led to the discovery of 361 proteins in N2a cells and 897 proteins in mouse brain neurons, identified as BIN1iso1-associated (proximal) or interacting proteins. A total of 92 proteins were common in both datasets, indicating that these are high-confidence BIN1- interacting or proximity proteins. SynapticGO analysis of the mouse brain dataset revealed that BIN1iso1-TurboID labeled 159 synaptic proteins, with 60 corresponding to the synaptic vesicle cycle. Based on phosphorylation site analysis of the neuronal BIN1iso1-TID interactome and related kinase prediction, we selected AAK1, CDK16, SYNJ1, PP2BA, and RANG for validation through immunostaining and proximity ligation assays as members of the BIN1 interactome in the mouse brain. By identifying several previously unknown proximal and potential interacting proteins of BIN1, this study establishes a foundation for further investigations into the function of neuronal BIN1.

## Introduction

The *Bridging Integrator 1* (*BIN1)* gene was identified in genome-wide association studies (GWAS) as the second most significant susceptibility locus for late-onset Alzheimer’s Disease (LOAD)^1–5^. BIN1 is a member of the BAR (BIN1/amphiphysin/RVS167) domain superfamily^6–8^ and is comprised of 20 exons that undergo alternative splicing to yield over a dozen BIN1 isoforms with distinct tissue and cell-type specific distributions^9^. BIN1 participates in several biological processes in cells, including endocytosis, cytoskeletal regulation, membrane dynamics, DNA repair, and apoptosis^9^. In peripheral tissue, altered splicing or aberrant expression of *BIN1* has been linked to cancer progression and centronuclear myopathy, and plasma BIN1 levels are associated with ventricular cardiomyopathy^9,10^. The prominent Alzheimer’s disease (AD)- associated single nucleotide polymorphisms (SNPs) are located in the *BIN1* 5’ region, 25-30 kb upstream of the coding sequence^2,11^, indicating that they reside in regulatory or enhancer regions that can confer AD risk by modifying BIN1 expression in brain cells^12,13^.

In the brain, at least 12 BIN1 isoforms are expressed to varying degrees in neurons, oligodendrocytes, and microglia^14–17^. The largest BIN1 isoform (BIN1iso1) is exclusively expressed in neurons; notably, a decrease in BIN1iso1 levels correlates with a significant loss of neurons observed in postmortem tissue from patients with AD^14^. In addition, an increase in ubiquitously expressed BIN1 isoform 9 (BIN1iso9) was observed in AD compared to healthy controls^14,15,18–20^. Neuronal BIN1 has four defined protein domains: the BAR domain, the CLAP (Clathrin-AP-binding) domain, the MBD (MYC-binding domain), and the SH3 (SRC homology 3) domain. The BAR, MBD, and SH3 domains are shared among other BIN1 isoforms, whereas the CLAP domain is specific to neuronal BIN1. The BAR domain enables BIN1 to associate with membranes, which is crucial for BIN1’s pivotal role in sensing and imposing membrane curvature and is essential in skeletal muscle for forming and maintaining T-tubule organization and structure to achieve proper excitation-contraction coupling^9,21,22^. The BIN1 MBD interacts with the c-MYC transcription factor, functioning as a tumor suppressor, with BIN1 protein levels altered in several cancer types and correlated with increased cancer metastasis^9^. The versatile SH3 domain binds to dynamin-1 and many other proteins with proline-rich domains (PRD), whereas the CLAP domain of neuronal BIN1 binds to clathrin and alpha adaptin-2 (within the AP-2 complex), as its name suggests^23–26^. BIN1’s ability to bind to dynamin-1, clathrin, and AP-2 implicates BIN1 in clathrin-mediated endocytosis. Moreover, early studies suggested a role for BIN1 in endocytosis, based on the observation that the SH3 domain of BIN1’s homolog, amphiphysin 1, efficiently inhibited synaptic vesicle endocytosis and receptor-mediated endocytosis^27,28^. Follow-up studies observed similar results with BIN1’s SH3 domain discovering that it binds to the same PRD on dynamin1 as amphiphysin 1 and similarly inhibits endocytosis by disrupting dynamin ring formation^29^. Indeed, temporal binding and clathrin-mediated endocytosis experiments conducted in NIH-3T3 mouse embryonic fibroblasts demonstrated that BIN1 plays a role in endocytosis and is recruited to clathrin-coated structures along with amphiphysin 1, endophilin 2, synaptojanin2β, and dynamin 1 immediately prior to membrane scission^30^. More recently, BIN1 has been suggested to contribute (through the recruitment of the microtubule motor dynein) to fast endophilin-mediated endocytosis^31^. Furthermore, BIN1 has also been proposed to be involved in EHD1-regulated endocytic recycling^32^. Much evidence supporting BIN1’s endocytic roles and protein interactions has been gathered from non-neuronal cells that do not replicate unique neuronal properties, such as rapid synaptic vesicle dynamics or the specialized protein landscape of neuronal synapses where BIN1iso1 natively functions.

In previous studies, we generated conditional knock-out mice lacking BIN1 expression in neurons to gain additional insight into neuronal BIN1 function. We found that the loss of BIN1 in excitatory neurons led to deficits in spatial learning and memory^17^. A 3D immuno-electron microscopy reconstruction analysis revealed that BIN1 localizes to the synapse, with a preference for localization at pre-synaptic terminals. The loss of BIN1 in excitatory neurons significantly increased the number of docked synaptic vesicles and the reserve pool, suggesting altered synaptic vesicle dynamics^17^. On the other hand, increasing neuronal BIN1iso1 expression in primary rat hippocampal neurons induced network hyperexcitability and heightened neuronal activity, which correlated with BIN1 interactions with L-type calcium channels, facilitated by tau^33^. Therefore, BIN1 plays a fundamental role in synaptic physiology in neurons. However, very little is known about how BIN1 functions at the synapse and the proteins that may influence BIN1 function(s) through dynamic interactions with it.

BIN1’s specific contributions to AD pathophysiology remain enigmatic. However, there is initial evidence that BIN1 can interact with proteins encoded by other GWAS candidate genes implicated in AD, such as CLUS^34^ and RIN3^35^. Of particular interest to the AD field, BIN1’s SH3 domain has also been shown to interact with tau *in vitro*^36–38^. Still, this interaction has not been reproduced *in vivo* under physiological conditions, and the significance of the binding between BIN1 and tau for AD pathogenesis is not fully understood^39,40^. The aforementioned studies have established a foundational understanding of BIN1’s domains and attributed functions based on robust interactions with fewer than a dozen proteins. Thus, significant gaps remain in our understanding of the diversity of BIN1 protein interactions and transient associations relevant to neuronal function. To address this gap, the present study sought to establish the interactome of human neuronal BINiso1 under normal, homeostatic conditions with the expectation that identifying proximal and interacting partners will collectively help define BIN1’s function in neurons. We employed a tagging and proximity labeling approach using the biotin ligase TurboID, which can biotinylate all proximal proteins within ∼20-nm radius in as little as 10 minutes^41,42^. TurboID and its predecessors, coupled with quantitative mass spectrometry, have been utilized to elucidate cell-type and subcellular-specific proteomes^41^. The improved kinetics of TurboID compared to its predecessors and the lack of toxicity relative to other peroxidase proximity ligases make it an appealing candidate for *in vivo* proximity labeling and protein-protein interaction (PPI) studies. This approach has already begun to revolutionize neuroscience, and its adoption is increasing at a rapid pace^43–46^.

Here, we describe the implementation of our approach to label BIN1 interacting and proximity proteins in cultured mouse neuroblastoma (N2a) cells and adult mouse brain neurons *in vivo*, followed by label-free quantification using mass spectrometry-based proteomics. BIN1iso1-TurboID fusion protein-mediated biotin labeling and proteomics identification resulted in the discovery of 361 proteins in N2a cells and 897 proteins in mouse brain neurons labeled as BIN1-associated (proximal) or interacting proteins. The two datasets shared 92 proteins, including those with known functions in endocytosis, as well as an unexpected number of mitochondrial proteins and nuclear histone modifiers. These shared BIN1 interactome proteins, along with the identification of phosphorylation sites and kinase predictions, enabled us to prioritize interactors for validation. We confirmed AAK1, CDK16, SYNJ1, PP2BA, and RANG as *bona fide* BIN1 proximal proteins in neurons within the brain. The *in vitro* and *in vivo* datasets we generated will provide a foundation for future investigations of neuronal BIN1 function and further clarify the regulatory protein network in which BIN1 plays its role as a risk factor for developing LOAD.

## Experimental Procedures Animals

All experimental procedures related to animal care and experimental manipulation were carried out in agreement with the Institutional Animal Care and Use Committee policies at the University of South Florida, Tampa. Mice were housed at 22 ± 2°C under a 12 h light/dark cycle with *ad libitum* access to food and water. *Bin1^fl/fl^* animals were generously provided by Dr. George C. Prendergast (Lankenau Institute for Medical Research)^47^. *Emx1*-IRES-*Cre* (JAX stock #005628) lines were obtained from The Jackson Laboratory (Bar Harbor, ME). We crossed the *Bin1^fl/fl^* strain with the *Emx1*-IRES-*Cre* driver lines to generate *Emx*-Cre:*Bin1* knockout mice (*Emx*-Cre:*Bin1* KO) and *Emx*-Cre littermate controls. The C57BL/6J (JAX stock #000664) mice were obtained from The Jackson Laboratory (Bar Harbor, ME).

## Molecular Cloning and Recombinant Virus Production

The lentiviral expression plasmids were generated using Gateway cloning. The following plasmids were purchased from Addgene: pENTR1A (Addgene #17398), C1(1-29)-TurboID- V5_pCDNA3 (Addgene #107173), pLX304 (Addgene #25890), and V5-TurboID-NES_pCDNA3 (Addgene #107169). The human BIN1 isoform 1 coding sequence was fused via a flexible 13X GS linker sequence to TurboID-V5 and cloned into the Gateway entry vector, pENTR1A; this fusion protein is referred to as BIN1iso1-TID. The cytosolic control protein, termed Cyto-TID, was generated by cloning V5-TurboID-NES into pENTR1A. These ORFs were transferred into the Gateway lentiviral destination vector, pLX304, using Gateway LR recombination (Invitrogen). Positive clones were confirmed by sequencing.

Recombinant AAV vectors were designed to express cytosolic TurboID (AddGene, Plasmid #107169) or the human BIN1 isoform 1 linked to TurboID via a flexible GS linker from the human SYN1 promoter to drive expression in neurons. The recombinant AAV vectors were constructed and packaged with AAV-PHP.eB capsid by VectorBuilder.

## Generation of Stably Transduced N2a Pools and TurboID Labeling

The pLX304-BIN1iso1-13X-TurboID-V5 and pLX304-V5-TurboID-NES lentiviruses were packaged in HEK293T cells and viral particles were collected using Mission lentiviral packaging mix (Sigma) according to the manufacturer’s protocol. Mouse neuroblastoma (N2a) cells were cultured in complete Dulbecco’s modified Eagle’s medium (DMEM) supplemented with 10% v/v fetal bovine serum (FBS), 1% Pen-Strep antibiotic and 1x GlutaMAX (Gibco). They were maintained in a humidified incubator with 5% CO_2_ at 37°C. Cells were transduced with lentiviruses, and stably transduced pools were selected and maintained under blasticidin selection (10 μg/ml). The expression of Cyto-TID and BIN1iso1-TID was confirmed by immunoblot analysis. For TurboID-mediated proximity biotinylation in stably transduced N2a cells, 50 µM of biotin was added to the complete media, and the cells were incubated for 30 minutes at 37°C. The cells were immediately placed on ice and washed with ice-cold PBS prior to lysis in RIPA buffer (150 mM NaCl, 50 mM Tris-HCl pH 8.0, 1% IGEPAL CA-630, 0.5% sodium deoxycholate, 0.1% SDS) supplemented with 1x Halt Protease and Phosphatase Inhibitor Cocktail (Thermo Fisher Scientific) and 1x PMSF. The lysates were processed for proteomics as described below.

## Recombinant AAV Delivery, Biotin Supplementation, and Tissue Collection

Two-month-old mice were briefly anesthetized with isoflurane, and 1 x 10^12^ viral genomes were systemically injected into the retroorbital sinus for AAV transduction. The mice were given 4 weeks for adequate AAV-mediated transgene expression before being supplemented with exogenous biotin for TurboID labeling. Biotin (Sigma B450, 0.5 mg/ml, titrated into solution in H_2_O with NaOH and adjusted to pH 7) was provided to the mice in their home cage water bottle. The mice had unrestricted access to biotin water for 5 days, with fresh biotin water provided on day 3.

After exposure to biotin in the water for 5 days, the mice were anesthetized with isoflurane and immediately transcardially perfused with chilled phosphate-buffered saline (10 mM, pH 7.4). Their brains were removed and cut in the midsagittal plane into two hemibrains. One hemibrain was post-fixed in phosphate-buffered saline containing 4% paraformaldehyde for 24 h and reserved for immunostaining. The other hemibrain was carefully dissected into hippocampus-cortex, midbrain-hindbrain and cerebellum, snap-frozen, and stored at −80°C for proteomics and other biochemical analyses.

## Immunofluorescence Staining

Post-fixed hemibrains were processed and embedded in paraffin blocks. Five-μm sections were cut, mounted, and deparaffinized in xylene, followed by rehydration through a decreasing ethanol dilution series. For epitope retrieval, the sections were incubated with Reveal Decloaker solution at 95°C for 30 min in the Decloaking Chamber NxGen, cooled to room temperature, and rinsed with tris-buffered saline (TBS). Slides were blocked for 1 h using Background Punisher. Da Vinci Green antibody diluent was used to dilute primary antibodies against V5 (Thermo Fisher #R96025), BIN1 (Proteintech #14647-1-AP), mAb BIN1 (clone 19H3)^40^, CDK16 (Proteintech #10102-1-AP), PP2Ba (Santa Cruz # sc-17808), SYNJ1 (Atlas Antibodies #HPS011916), AAK1 (Atlas Antibodies #HPA020289), RANG (Atlas Antibodies #HPA065868), and Streptavidin conjugates (Invitrogen #84547). Primary antibodies were incubated overnight at 4°C while secondary antibodies were incubated for 2 h at room temperature. Nuclei were stained using Hoechst 33342 before mounting coverslips with VectaShield mounting media and sealing with nail polish.

## Proximity Ligation Assay

Paraffin sections were used for PLA assays, and the immunofluorescence processing steps outlined above were followed up to the blocking step. During this step, the Duolink PLA manufacturer protocol and reagents were used (Millipore Sigma). Primary antibodies (listed above) were diluted in Duolink PLA antibody diluent and incubated overnight at the concentrations recommended by the manufacturer. The following probes were used to bind the primary antibody: Duolink *In Situ* PLA Probe Anti-Mouse MINUS and Duolink *In Situ* PLA Probe Anti-Rabbit PLUS with the FarRed detection reagent to detect and visualize the proximal protein-protein interactions.

## Image Acquisition and Quantification

Images were acquired on an automated Nikon Eclipse Ti2 microscope fitted with the Yokogawa spinning disk field scanning confocal system and Photometrics PRIME 95B sCMOS camera, using 4x, 20x, 60x, and 100X objectives. High-magnification z-stack images were deconvolved using Huygens software (24.10). Spearman colocalization analysis and colocalization object maps (Synaptophysin [SYP] or PSD95 and streptavidin) were quantified using Huygens Colocalization Analyzer. Total presynaptic (SYP positive objects) and postsynaptic (PSD95 positive objects) counts were generated from z-stacks using Huygens Object Analyzer. The ratio was quantified from the total number of colocalized objects (synapses positive for biotinylated proteins) / the total number of synapses. Images were processed and converted into 2D projections with Fiji/ImageJ software.

## Immunoblot Analysis

Cortex and hippocampus were dissected from the mouse brain and homogenized in 10% weight/volume RIPA lysis buffer (150 mM NaCl, 50 mM Tris-HCl pH 8.0, 1% IGEPAL CA- 630, 0.5% sodium deoxycholate, 0.1% SDS) supplemented with 1x Halt Protease and Phosphatase Inhibitor Cocktail (Thermo Fisher Scientific) and 1x PMSF. DNA was sheared using a probe sonicator. Proteins were extracted from N2a cells by trituration in RIPA lysis buffer. Aliquots of protein samples were run on 4-20% Bis-Tris gels and the blots were probed with primary antibodies against V5 (Thermo Fisher #R96025), BIN1 (Proteintech #14647-1-AP) and biotinylated proteins with streptavidin conjugated to infrared dye-IR680 (LI-COR #926- 68076) or IR800 (LI-COR #926-32230). The blots were developed with infrared dye-conjugated IR680 (LI-COR #926-68070) and IR800 (LI-COR #926-32213) secondary antibodies and imaged using the Odyssey Infrared Imaging System (Li-COR Biosciences). Following the elution of captured biotinylated proteins from streptavidin magnetic beads, samples were checked for total protein using Coomassie staining. Gels were fixed in 50% methanol and 10% glacial acetic acid for 1 h at room temperature in a sealed container on a rocking incubator. The gels were stained for 30 min (0.1% Coomassie Brilliant Blue R-250, 50% methanol, 10% glacial acetic acid) and then destained with a destaining solution (40% methanol and 10% glacial acetic acid).

## Protein Processing for Mass Spectrometry

The streptavidin pull-down method for precipitating biotinylated proteins from total protein lysates was adapted from a published protocol with minor modifications^42^. In brief, 1 mg of total protein from N2a cell lysates was incubated with 55 µl of Pierce streptavidin magnetic beads (ThermoFisher Scientific); whereas 2.5 mg of total protein from hippocampal-cortex lysates was incubated with 100 µl of Pierce streptavidin magnetic beads. The total protein lysates were incubated overnight at 4°C. The following day, the beads with bound proteins were washed in the following series of solutions: RIPA buffer, 1 M KCl, 0.1 M Na_2_CO_3_, 2 M urea in 10 mM Tris-HCl pH 8.0, RIPA buffer, and PBS. The washed beads with bound proteins were resuspended in 50 mM ammonium bicarbonate with 5% SDS, 50 mM DTT, and 5 mM biotin.

Proteins were eluted from the beads using a thermal mixer set to 98°C and 1000 RPM for 10 mins. The eluted proteins were reduced with 70 mM DTT for 10 mins, alkylated with 140 mM IAA for 30 mins, and loaded onto an S-Trap micro spin column (Protifi) according to manufacturer’s protocol. After sample clean-up, the trapped proteins were digested with 1 µg of Trypsin (Promega) and the resulting peptides were eluted in a series of buffers including, 50 mM ammonium bicarbonate, 0.2% formic acid in LC-MS grade H_2_O, and 50% acetonitrile containing 0.2% formic acid, all collected into one final Eppendorf tube. The eluted samples were then diluted in 0.1% formic acid to reduce acetonitrile concentration, desalted in C18 SPE columns (Waters) with several washes of 0.1% formic acid, eluted in 50% acetonitrile containing 0.1% formic acid, and dried down using a speed vacuum manifold with a cold vapor trap. Dried peptides were resuspended in 0.1% formic acid in water and transferred to autosampler vials for LC-MS/MS analysis.

## Mass Spectrometry-based Proteomic Analysis

Resuspended samples were injected to achieve a normalized on-column amount (based on total ion chromatogram signal) and then separated via ultra-high performance liquid chromatography (UHPLC) nanoElute (Bruker) using an Aurora Ultimate CSI UHPLC reversed-phase C18 column (25 cm x 75 µm i.d., 1.7 µm C18, IonOpticks) heated with a column oven set to 50°C. The column was in line with a trapped ion mobility-QTOF mass spectrometer, the timsTOF Pro (Bruker), and the majority of peptides were eluted with mobile phases A (0.1% formic acid in water) and B (0.1% formic acid in acetonitrile) using a 90 min gradient of 2-25% B. The total runtime of 120 min included an additional ramp-up to 37-80% B to clean the column followed by a blank injection prior to analysis of each experimental sample. DIA-PASEF scan mode was utilized with mass widths of 25 Da (mass windows with no overlap), one mobility window of 0.7-1.4 1/K0 [V·s/cm^2^] spanning 250-1425 m/z, 1.48 s estimated cycle time, and set with a collision energy of 20 eV for a base of 0.60 1/K_0_ [V·s/cm^2^] and 59 eV for a base of 1.60 1/K_0_ [V·s/cm^2^]. Ion mobility and m/z calibration were performed using three calibrant ions at 622, 922, and 1222 m/z (Agilent). DIA data for both the N2a samples and hippocampal-cortex tissue samples were searched separately in DIA-NN (v. 1.8.1) using identical default settings. A predicted library was generated from the Uniprot *Mus musculus* database (UP000005640, 55,315 entries) with the manual addition of streptavidin as an additional contamination. Within the label-free quantification (LFQ), the match-between-runs (MBR) feature was implemented using an FDR cutoff of 1%. The following settings were also selected: both mass accuracies at 15 ppm, single pass mode, genes as the protein inference, robust LC (high precision), RT-dependent cross-run normalization, and smart profiling. A subset of the DIA data obtained from the male hippocampal-cortex tissue samples was re-searched with the purpose of identifying phosphorylated peptides in DIA-NN (v.1.8.1) using the same settings but with an alternative predicted library that consisted of phosphorylation of STY as a variable modification, with the maximum number of variable modifications set to one.

## Statistical Analysis of LC-MS/MS Data

Statistical analysis of the N2a interactome dataset was performed as previously described^48^, with updates and minor changes to accommodate specific settings for this data set within Perseus (v. 2.0.10.0). In summary, LFQ intensity values were annotated, log2 transformed, and protein groups that did not meet identification/quantitation in 100% of at least one of the two experimental groups (Bin1iso1-TID and Cyto-TID) were filtered out to achieve the highest confidence due to the number of replicates used in this comparison (n=3). The remaining protein groups with missing values had those values replaced with the imputation function using the normal distribution option, set with a width of 0.5 and a downshift of 1.6 to fall within the lower abundance area of the curve^49^. A Welch’s T-test was applied with a p-value cutoff of < 0.05, and proteins that did not pass this threshold were filtered out before normalizing for hierarchical clustering. Gene ontology (GO) annotations related to GOBP, GOMF, GOCC, and KEGG pathways were then added using the *Mus musculus* Gene Ontology (GO) reference set and the resulting list was exported for further analysis in Excel. An additional z-score cutoff of >1 was also implemented to the previously applied p-value cutoff^50^. Protein groups that met the p-value cutoff, z-score cutoff, and were positive in the t-test difference were considered significantly enriched proteins representing the Bin1 interactome. Hippocampal-cortex tissue samples were analyzed similarly, without differentiating between male and female samples, and all of the Bin1iso1-TID group (n=11) was compared to the control Cyto-TID group (n=9) after filtering values based on 75% identification/quantification in at least one group. Imputation for missing values was set with a width of 0.4 and a downshift of 1.7. The subsequent Welch’s t-test, hierarchical clustering, z-score, and t-test difference steps were identical to those performed for the N2a dataset above.

## Bioinformatics and Data Visualization

Functional protein-protein interaction networks were generated using the STRING database (Version 12.0)^51^, using all evidence sources in conjunction with Cytoscape (Version 3.10.3) for customized network visualization and annotation. The details on edge score threshold and z-score cutoff for proteins visualized in each network are listed under the figure legends. A gene ontology (GO) analysis for GOBP, GOMF, and GOCC was conducted for positive protein hits from each dataset with a z-score >1 using ShinyGO (Version 0.80, based on Ensembl Release 104, archived on Oct 25, 2024)^52^, with a default background, and an FDR < 0.05. P-values were calculated using the hypergeometric test, and FDRs were computed via the Benjamini-Hochberg method to correct for multiple testing. Fold enrichment is defined as the percentage of genes in the z>1 hits that are in a pathway divided by the corresponding percentage in the background genes. While FDR measures statistical significance, fold enrichment indicates effect size. Synapse GO (SynGO release 1.2)^53^ was used to analyze positive protein hits with z-score >1 from the BIN1 mouse brain neuron dataset for their GOBP and GOCC, using default background set of all brain-expressed genes, with an FDR < 0.05. For each ontology term, a one-sided Fisher exact test was performed, followed by multiple testing correction using FDR. GraphPad Prism (Version 10.1.2) was utilized for graphical presentation. BioRender.com was used to create schematics.

## Experimental Design and Statistical Rationale

The sample conditions for *in vitro* analysis were prepared as follows: cell type (neuroblastoma N2a) and TurboID fusion protein (Cytosolic-TurboID or BIN1iso1-TurboID).

Each sample condition included three biological replicates. For the *in vivo* experiment, the sample conditions were C57BL6J mice and the TurboID fusion protein (Cytosolic-TurboID or BIN1iso1-TurboID). The *in vivo* experimental design was based on the preliminary proteomic analysis from *in vitro* studies, utilizing the following rationale and power analysis. Considering the potentially higher variability *in vivo* (average technical replicate precision of CV<5%^54^ with average biological replicate precision *in vivo* typically at CV<20% across ∼5K proteins), a group size of n=6 will be suffice to achieve a power of 0.90, given α=0.05, 30% CV, and >2-fold enrichment (the two-tailed difference between two independent means as calculated by G*Power 3.1.9.7). Nine biological replicates were analyzed in the Cyto-TurboID group, whereas 11 biological replicates were analyzed in the BIN1iso1-TurboID group. Consequently, a Welch’s t-test with a p-value cutoff of < 0.05 was chosen for the two independent samples. A z-score cutoff of > 1 was also applied, which has been shown to control FDR with minimal impact on sensitivity for LFQ-based proteomics^50^. Statistical analysis was performed as described above.

## Results

### TurboID fused to human BIN1iso1 successfully localizes in the cytoplasm and efficiently biotinylates proximal proteins

TurboID, an optimized biotin ligase, successfully labels proximal proteins within a ∼10- 20-nm radius in as little as 10 min of exposure to biotin in live cells^40^. We designed a BIN1- TurboID fusion protein based on human BIN1iso1 to identify the neuronal BIN1 interactome. In order to maintain the integrity of the N-terminal BAR domain, which is essential for association with cellular membranes, we fused TurboID and a V5-epitope tag to the C-terminus of BIN1 using a flexible GS linker in between **(Fig. 1A)**. Since BIN1 is a cytosolic adaptor protein, we opted for TurboID with a nuclear export sequence (Cyto-TID) that would freely biotinylate proteins in the cytosol as our negative control. We generated pools of mouse neuroblastoma (N2a) cells stably transduced with lentiviruses encoding the fusion proteins. First, we confirmed the expression of BIN1iso1-TID and Cyto-TID and characterized intracellular proximity biotinylation mediated by TurboID **(Fig. 1B)**. Increasing concentrations of exogenous biotin (0, 25 µM, and 50 µM) were added to the cells for 10 mins of labeling to assess the background activity of the TurboID fusion proteins and determine the optimal dose for subsequent experiments **(Fig. 1C)**. Immunoblots of the lysates were probed with anti-V5 to visualize the TurboID fusion proteins and with streptavidin to detect all biotinylated proteins. In the samples without exogenous biotin, there was low-level TurboID activity due to normal biotin supplementation in the cell culture media, along with several bands common to all lanes that correspond to endogenous biotinylated proteins, mostly carboxylases (72, 75, 130 kDa)^42,55^. The addition of exogenous biotin significantly increased streptavidin signal intensities in a dose-dependent manner in both the Cyto-TID and BIN1-TID groups, as well as some self-biotinylation, as indicated by the overlap of V5-epitope and streptavidin signals. Incubating cells with 50 µM of exogenous biotin produced a stronger signal relative to background labeling; this concentration was used for subsequent experiments. Next, exogenous biotin (50 µM) was added to the cells and incubated for 10, 20, 30, and 60 min before cell lysis and analysis to determine the optimal labeling time. Immunoblots of the lysates were analyzed using anti-BIN1 and V5 antibodies to visualize endogenous BIN1 and V5-tagged BIN1iso1-TID. The immunoblot displayed three prominent endogenous mouse BIN1 isoforms at 50-65 kD and BIN1iso1-TID at 100 kD. The V5 antibody labeled ∼100 kD BIN1iso1-TID at 100 kD and ∼35 kD Cyto-TID as expected (**Fig. 1D**). The blot also indicated modest BIN1iso1-TID expression relative to endogenous BIN1 and comparable expression of both TurboID proteins, allowing for biotinylation of proximal proteins without encountering overexpression artifacts. Further analysis of the lysates revealed robust biotinylation at all time points, as demonstrated by streptavidin detection, reaching a peak level of biotinylation at 30 min **(Fig. 1E)**. We utilized this labeling time for subsequent experiments. After optimizing our biotinylation conditions, we performed immunofluorescence labeling to determine the localization of BIN1iso1-TID in N2a cells compared to endogenous mouse BIN1. We also assessed the extent of overlap between the BIN1iso1-TID or Cyto-TID and biotinylated proteins detectable by streptavidin, which was anticipated to be relatively high. Using confocal microscopy, we observed the cytosolic distribution and punctate localization of BIN1iso1-TID and Cyto-TID, as indicated by V5 immunostaining **(Fig. 1F)**. In N2a cells expressing BIN1iso1-TID, there was substantial overlap between V5 and anti-BIN1 immunofluorescence, indicating that BIN1iso1-TID has a distribution similar to that of endogenous mouse BIN1. Finally, detection by streptavidin revealed a similar distribution of biotinylated proteins throughout the cytosol in both groups, suggesting that Cyto- TID serves as an effective spatial reference control for distinguishing the BIN1iso1-specific interactome through enrichment analysis.

**Fig 1.**
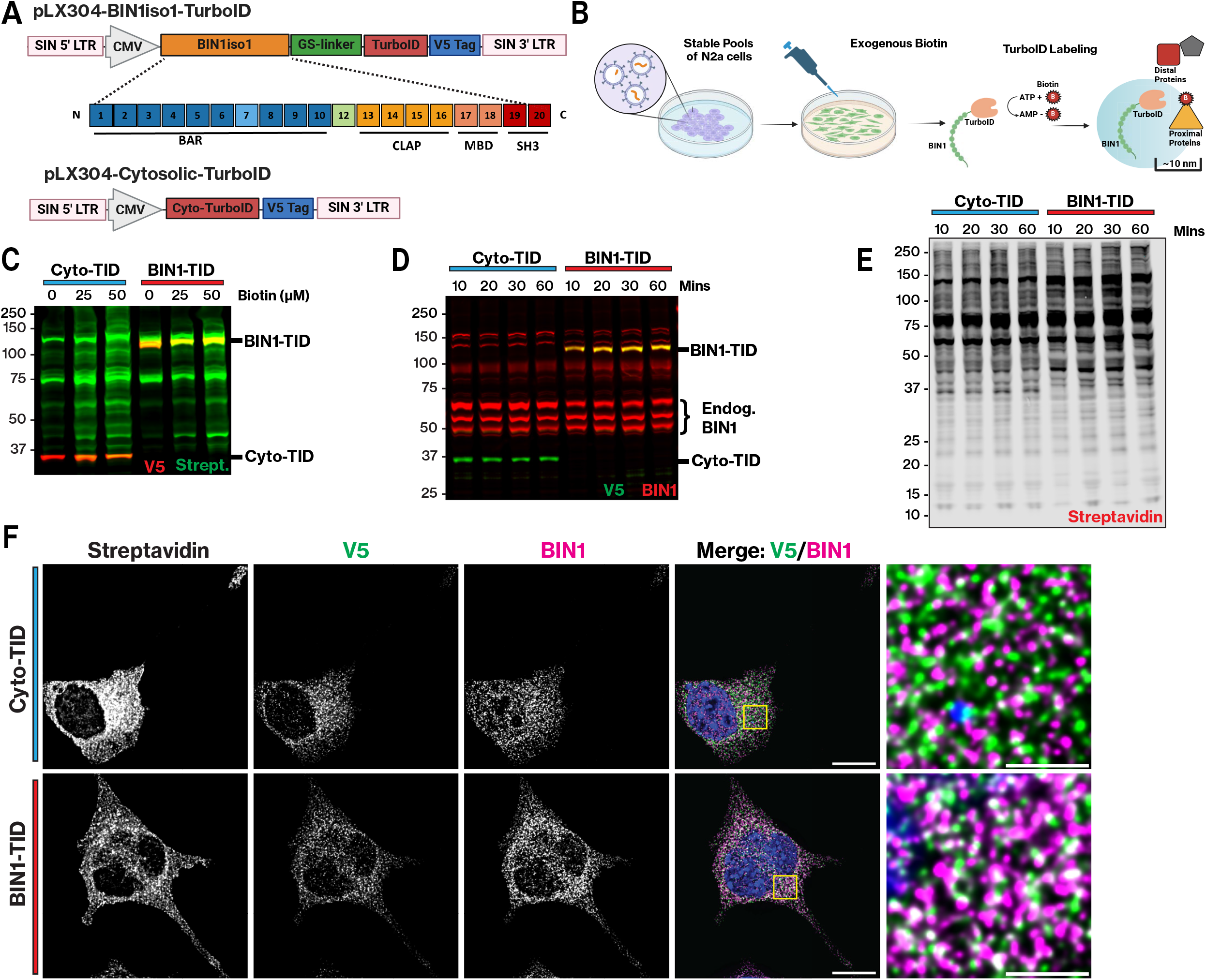
N2a cells stably expressing BIN1iso1-TID and Cyto-TID localize to the cytosol and biotinylate proximal proteins. **A)** Schematic of lentiviral vectors expressing Human BIN1iso1 fused to TurboID via a flexible GS linker and a V5-epitope tag and Cytosolic-TurboID with an NES and a V5-epitope tag. **B)** Schematic of experimental design. Stable pools of neuroblastoma (N2a) cells expressing BIN1iso1-TID were provided exogenous biotin and incubated for the times mentioned below prior to washing and lysis. **C)** Biotin dose-response (- biotin, 25 µM, 50 µM). Exogenous biotin was added to the cells for 10 minutes of labeling prior to washing and lysis. Western blot shows anti-V5 (680) and biotinylated proteins (streptavidin- 800). **D)** 50 µM of exogenous biotin was added for 10, 20, 30, and 60 mins. Western blot is probed with anti-BIN1 (680) and anti-V5 (800), revealing endogenous BIN1 and expression of TurboID fusion proteins. **E)** Same lysates as above run on a blot and probed with streptavidin, detecting quantity of biotinylated proteins through 60 min time course. **F)** Immunofluorescence (IF) of N2a stable cells expressing BIN1iso1-TID and Cyto-TID fusion proteins. Anti-V5, anti- BIN1, streptavidin, and Hoechst. Images are deconvolved z-stacks displayed as a sum projection (scale bar 10 µM). The boxed region in the V5 and BIN1 image overlay is shown at a higher magnification on the right.

## BIN1iso1-TID identifies both known and unknown proximal proteins in mouse N2a neuroblastoma cells

We incubated stable N2a pools expressing BIN1iso1-TID or Cyto-TID with 50 µM biotin for 30 min and processed the cell lysates for mass spectrometry. In brief, biotinylated proteins were captured using streptavidin magnetic beads and then eluted from the beads. The eluate was run on a gel and stained with Coomassie to check for the quality of the available proteins and to ensure that both groups had sufficient protein for downstream mass spectrometry **(Supplemental Fig. 1A)**. Both groups had comparable amounts of protein, which were then processed using S-Trap^56–58^, followed by trypsin digestion and desalting **(Fig. 2A)**. We utilized data-independent acquisition (DIA) mass spectrometry to acquire the data because this approach has demonstrated increased depth of coverage, reproducibility, and sensitivity compared to data-dependent acquisition^59,60^, making it optimal for discovery research. DIA mass spectrometry successfully and reproducibly identified ∼6,000 and ∼5,200 unique proteins (FDR, 0.01) captured by streptavidin from stable N2a cells expressing Cyto-TID and BIN1iso1-TID, respectively **(Fig. 2C)**. We implemented a stringent data analysis approach **(Fig. 2B)** to delineate biotinylated proteins that are more abundant in the BIN1iso1 interactome when compared to proteins labeled by Cyto-TID. In brief, positive hits were based on Welch’s T-test p-value < 0.05, filtered based on a coefficient of variation (CV) < 30% for the BIN1 group and a z-score cutoff >1. For each protein, the z-scores reflect the difference between the protein fold change and the mean of the population fold changes relative to the population standard deviation (see Experimental Procedures)^50^. This analysis identified a set of 361 proteins enriched in the N2a BIN1iso1-TID samples with a z-score >1. This dataset represents the BIN1iso1 interactome, including proximal proteins and direct BIN1iso1 interactors. We used the Search Tool for the Retrieval of Interacting Genes/Proteins (STRING) to generate a protein-protein interaction (PPI) network of the 114 enriched proteins with a z-score >2, as these proteins are more likely to be direct interactors and allow for easier visualization of the PPI network **(Fig. 2D)**. The most highly enriched protein in the BIN1iso1-TID interactome was endogenous mouse BIN1 **(Fig. 2E and Fig. 2F)**, highlighted in yellow in cluster 3. This finding is not surprising, as it is well-established that BIN1 forms homodimers^26^. BIN1’s immediate neighbors directly connected to BIN1 in cluster 3 include SYNJ1, SYNJ2, and LIN54. Both SYNJ1 and SYNJ2 are involved in clathrin-mediated endocytosis and phosphatidylinositol signaling, consistent with previous research demonstrating BIN1’s interaction with these proteins^61^ and several others, which are annotated in red on the volcano plot **(Supplementary** Fig.1B)^9^.

**Fig 2.**
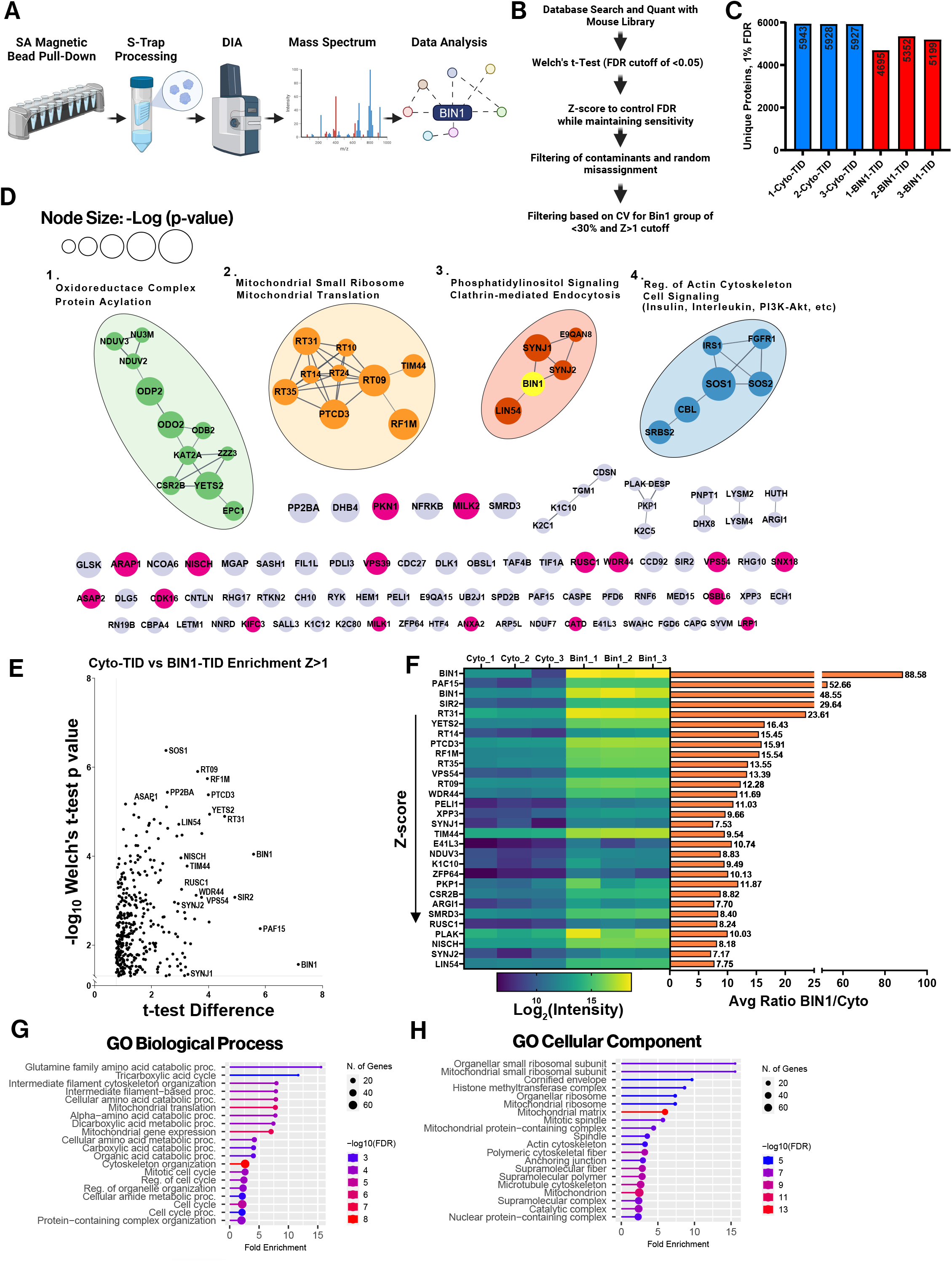
BIN1iso1-TurboID interactome in N2a cells identifies known and unknown BIN1- proximal or interacting proteins. **A)** Overview schematic of sample processing for mass spectrometry. **B)** Data analysis pipeline for mass spectrometry from database search and quantitation through final cutoffs (CV<30%, Z-score >1). **C)** Bar graph displaying unique proteins (detection FDR<0.01) for N2a BIN1iso1-TID and Cyto-TID samples in triplicate (n=3). **D)** The Search Tool for the Retrieval of Interacting Genes/Proteins (STRING) network generated for N2a BIN1iso1-TID interactome samples for Z-score >2.The edges connecting nodes in a STRING network are determined by scores that indicate the confidence level regarding the likelihood that a PPI is valid, based on available evidence, ranging from 0 to 1. A score of 0.5 suggests that every second interaction could be a false positive. Thus, we opted to filter the edges connecting nodes in the network to a high confidence score of > 0.7. These edges signify a predicted functional relationship and/or physical interaction, while the size of the nodes corresponds to the -Log (Welch’s T-test p-value). Four major clusters annotated and labeled according to representative terms. Singlet nodes in magenta correspond to cytoskeleton regulation, endocytosis, and vesicle-mediated transport. **E)** Volcano plot displaying T-test Difference vs. -Log10(Welch’s T-test p-value) for N2a BIN1iso1-TID interactome (Z-score >1) F) On the left, heat map of Log2(intensity) for top 30 proteins, by descending Z-score and a horizontal bar graph on the right displaying the average ratio BIN1iso1-TID/Cyto-TID. **F)** Lollipop graph of Gene Ontology (GO) term analysis for biological process. Lines and lollipops are colored according to -Log10(FDR) and the lollipop sizes correspond to the number of genes. **G)** GO term analysis for cellular component. Lines and lollipops are colored according to - Log10(FDR) and the lollipop sizes correspond to the number of genes.

While proteins involved in clathrin-mediated endocytosis are expected to be represented in the BIN1 interactome, BIN1 proximity labeling also identified many unexpected proteins, suggesting that BIN1 has a complex role in cellular biology. Cluster 1 includes several proteins typically associated with the nucleoplasm, protein acylation, and histone modification (KAT2A, ZZZ3, CSR2B, YETS2, EPC1). YETS2, which contains several proline-rich motifs predicted to enable SH3 domain binding, was one of the most highly enriched proteins in the mouse neuroblastoma dataset, with an average ratio of 16.4-fold. YETS2 is part of the ADA Two A-Containing (ATAC) histone acetyltransferase complex involved in transcription regulation and has been observed to act specifically as a selective histone crotonylation reader, with site specificity for H3K27^62^. A recent study found that YETS2 colocalizes with H3K27ac and controls a transcriptional program essential for lung cancer tumorigenesis^63^. BIN1 has been extensively studied in cancer research and characterized as a tumor suppressor through its interactions with the MYC transcription factor^9^. However, MYC was not enriched in our N2a BIN1 interactome dataset. Two other prominent proteins in the interactome indicate that BIN1 may play a role in the cell cycle and DNA repair. LIN54 (7.75 average fold ratio) and PAF15 (52.6 average fold ratio) are involved in cell cycle regulation but perform differing roles. LIN54 is directly connected to mouse BIN1 within the network and is reported to be a member of the multi-subunit DP, RB-like, E2F, and MuvB (DREAM) complex, which is involved in the transcription activation or repression of cell cycle genes depending on the cell cycle phase^64^.

LIN54 possesses a DNA binding motif and has been found necessary for cell cycle progression^65^. Despite LIN54’s connection to mouse BIN1 in the network, there is very little direct biochemical evidence in the literature linking them directly, requiring follow-up studies.

One of the most surprising findings in our study is the labeling a set of mitochondrial proteins by BIN1iso1-TID; 5 of the top 10 most enriched proteins in the BIN1 interactome are localized to the mitochondria. These are represented in Cluster 1, which includes components of the oxidoreductase complex and aerobic respiration (NDUV3, NU3M, NDUV2, ODP2, and ODO2), as well as in Cluster 2 of the network, which features mitochondrial translation machinery (RT09, RT10, RT14, RT24, RT31, RT35, PTCD3, and RF1M). Indeed, a Gene Ontology (GO) term analysis shows that the BIN1 interactome is enriched with proteins involved in the biological process of mitochondrial translation **(Fig. 2G),** with specific enrichment for proteins in the mitochondrial small ribosomal subunit **(Fig. 2H)**. BIN1 has no known role in mitochondrial biology, despite many of these BIN1isoTID-labeled proteins localizing to the mitochondrial matrix or mitochondrial ribosomes.

Finally, there are numerous singlet nodes in the network. This underscores the current state of limited understanding regarding PPIs and indicates that many proteins lack sufficient evidence to connect their nodes with high confidence. However, many of these proteins have GO molecular functions such as cytoskeletal protein binding and SH3 domain binding **(Supplemental Fig. 1C)**. Indeed, BIN1 is reported to possess a proline-rich motif in the CLAP domain that facilitates binding with other SH3 domain proteins^36^. Some of these singlets, highlighted in magenta in the network, are consistent with a functional role for BIN1 in cytoskeleton regulation (MILK2, PKN1, ARAP1), endocytosis (SNX18), and vesicle-mediated transport (VPS39, VPS54, WDR44, ASAP2, CDK16)^9^. Despite the absence of high-confidence connections, some of the most highly enriched proteins in the N2a BIN1iso1 interactome are singlet nodes, such as PAF15, also referred to as PCNA-associated factor (PCLAF). PCLAF (52.6-fold enrichment in BIN1 N2a interactome), a PCNA binding protein linked to DNA repair and centrosome regulation, is overexpressed in neuroblastoma and has been shown to accelerate cell proliferation and cell cycle progression, and is associated with poor outcomes in cancer patients^66^. PCLAF is a small 15 kDa protein that harbors an SH3 binding motif, suggesting that BIN1 may directly interact through its SH3 domain. However, no formal connection between PCLAF and BIN1 is currently represented in the literature.

## Retro-orbital AAV delivery of BIN1iso1-TID effectively transduces neurons and biotinylates proximal proteins in the mouse brain

Whereas N2a cells served as a simpler *in vitro* model, we extend the studies above, by expressing BIN1iso1-TID in neurons within the mouse brain to characterize the BIN1 neuronal interactome *in vivo*. To achieve this, we designed a recombinant AAV and directed expression of human BIN1iso1-TID specifically in neurons using the human synapsin 1 (SYN1) promoter^67,68^. The recently developed AAV serotype capsid AAV-PHP.eB effectively targets the central nervous system (CNS) and transduces neurons with high efficiency following a systemic retro-orbital intravenous injection of 1 x 10^11^ vector genomes^69^. This approach is appealing due to its technical efficiency and noninvasive delivery of rAAV into the adult mouse brain compared to stereotaxic injections. We administered 1 x 10^12^ vector genomes through retro-orbital injections into 2-month-old mice and allowed four weeks for transgene expression. For *in vivo* proximity labeling, we supplemented biotin (0.5 mg/ml) in the drinking water for 5 days^70^ **(Fig. 3A)**. We harvested the mouse brains, dissecting the hippocampus and cortex from one hemi-brain for streptavidin enrichment and proteomics analysis. First, we determined that rAAV-mediated expression of Cyto-TID and BIN1iso1-TID in adult mouse brains led to comparable proximity biotinylation of proteins in the cortex and hippocampus **(Fig. 3B)**. We then examined sagittal sections from mice transduced with Cyto-TID and BIN1iso1-TID to assess the distribution of cells transduced by rAAV-PHP.eB through immunofluorescence staining and the overall localization of biotinylated proteins via streptavidin detection **(Fig. 3C)**. We observed expression of transgene-derived proteins in the olfactory bulb, midbrain, cerebellum, and brain stem. While the expression in the cortex and hippocampus was relatively lower, higher magnification images revealed readily detectable neuronal expression of Cyto-TID and BIN1iso1-TID, as well as biotinylated proteins in the cortex and hippocampus, visualized through V5 immunostaining and streptavidin detection, respectively (**Fig. 3D**). Consistent with the localization of endogenous BIN1 in neuronal synapses, the biotinylation mediated by BIN1iso1-TID is enriched in synaptic terminals, appearing as punctate staining. In contrast, biotinylation mediated by Cyto-TID is more abundant in the neuronal soma. Further immunostaining of the cortex using antibodies against BIN1, V5, and biotin demonstrates overlap between BIN1 and V5 in the cortex of the BIN1iso1-TID group, as expected **(Supplementary** Fig. 2A). In the hippocampus, antibodies against V5 and biotin also show overlap between BIN1iso1-TID and biotinylated proteins in the BIN1iso1-TID group, as anticipated **(Fig. 3E)**. A strong V5 signal is present in the CA2 pyramidal cells of the hippocampus, along with substantial streptavidin detection of biotinylated proteins following the expression of Cyto-TID and BIN1iso1-TID. Overall, these data indicate that BIN1iso1-TID is expressed in neurons in the adult mouse brain, and TurboID effectively biotinylates proteins for quantitative mass spectrometry analysis.

**Fig 3.**
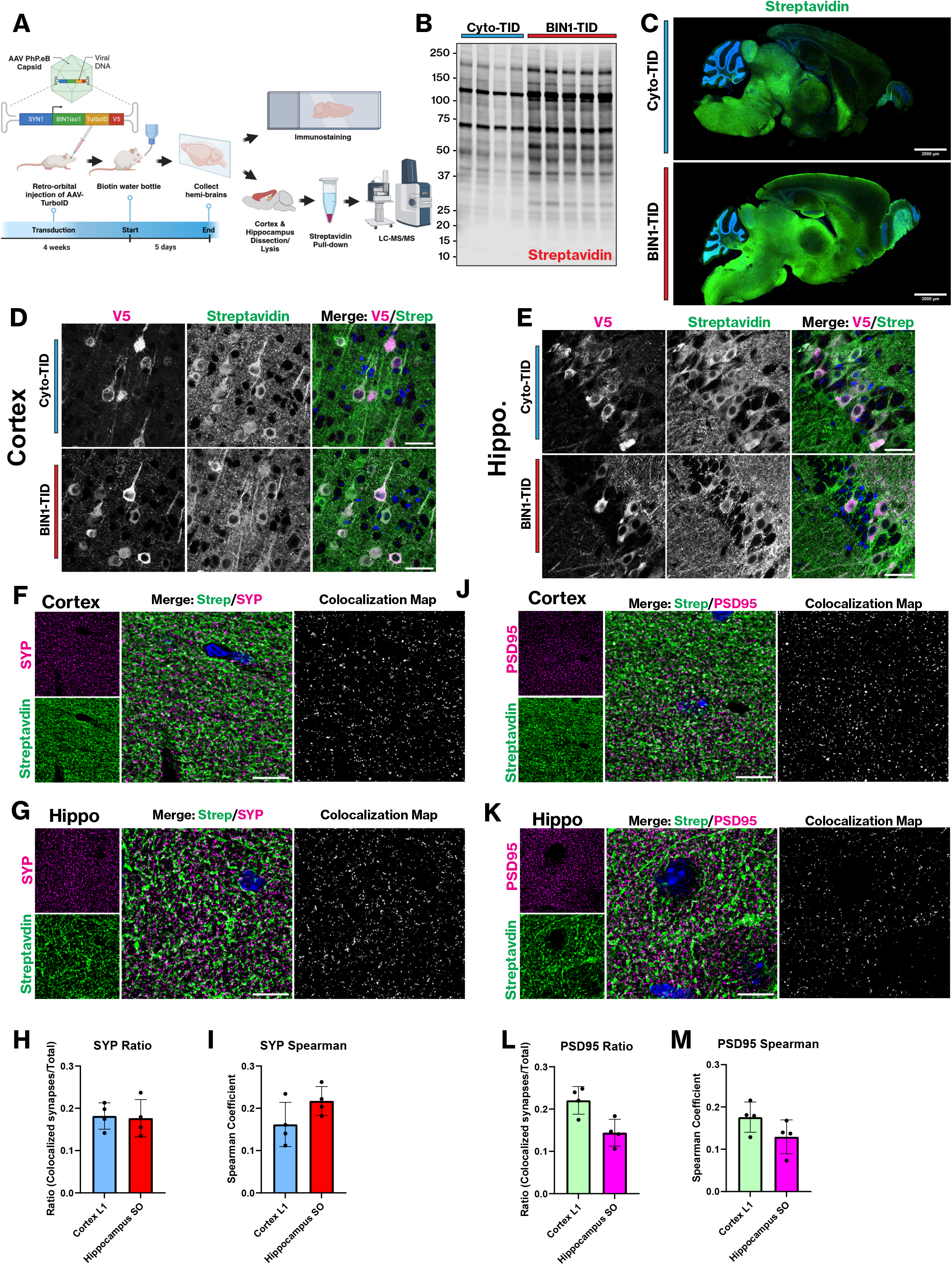
Retroorbital injection of rAAV human BIN1iso1-TID in mice efficiently transduces neurons and biotinylates proximal proteins in neurons. **A)** Schematic overview of experimental design for *in vivo* rAAV delivery. The rAAV uses a recently developed AAV serotype capsid AAV-PHP.eB with a human SYN1 promoter to drive expression of BIN1iso1-TID and Cyto-TID in neurons. A flexible GS linker fuse BIN1iso1 to TID with a V5 epitope tag. The AAV-TID was injected RO into 2-month-old mice and transduction was allowed for four weeks. Following this period, exogenous biotin (0.5 mg/ml) was supplemented in the mouse water for 5 days, followed by brain collection. Brains were split in half for immunofluorescence staining, and the hippocampus and cortex were dissected for quantitative mass spectrometry. Biotinylated proteins were pulled out with streptavidin beads and lysates were processed using S-Trap before LC-MS/MS. **B)** A Western blot using combined hippocampus and cortex lysates was probed with streptavidin (800) to detect biotinylated proteins. **C)** IF detection of biotinylated proteins via streptavidin. Sagittal sections from Cyto-TID and BIN1iso1-TID brains (Scale bar 2 mm). **D)** IF staining of BIN1-TID and Cyto-TID mouse hemi-brains with anti-BIN1, anti-V5, streptavidin, and Hoechst. Images are focused on the cortex and the merged image shows BIN1/V5 overlap. **E)** IF staining of BIN1-TID and Cyto-TID mouse hemi-brains with anti-BIN1, anti-V5, streptavidin, and Hoechst. Images are focused on the hippocampus and the merged image shows BIN1/V5 overlap. **F)** IF staining BIN1-TID mouse brains using anti-synaptophysin to label presynapses and streptavidin to detect biotinylated proteins. Images are deconvolved z-stacks of cortex projected as a sum. Colocalization analysis in Huygens generated the colocalization map (Scale bar 10 µm). **G)** IF staining BIN1-TID mouse brains using anti-synaptophysin to label presynapses and streptavidin to detect biotinylated proteins. Images are deconvolved z-stacks of hippocampus projected as a sum. Colocalization analysis in Huygens generated the colocalization map (Scale bar 10 µm). **H)** Quantification for the ratio of colocalization map SYP/streptavidin objects divided by the total SYP positive objects in the z-stack for cortex and hippocampus images (Huygens colocalization and object analysis, n=4 brains, 6 images per mouse per region). **I)** Quantification of Spearman colocalization coefficient for SYP/streptavidin hippocampus and cortex images (Huygens colocalization analysis, n=4 brains, 6 images per mouse per region). **J)** IF staining BIN1-TID mouse brains using anti- PSD95 to label postsynapses and streptavidin to detect biotinylated proteins. Images are deconvolved z-stacks of cortex projected as a sum. Colocalization analysis in Huygens generated the colocalization map (Scale bar 10 µm). **K)** IF staining BIN1-TID mouse brains using anti-PSD95 to label postsynapses and streptavidin to detect biotinylated proteins. Images are deconvolved z-stacks of hippocampus projected as a sum. Colocalization analysis in Huygens generated the colocalization map (Scale bar 10 µm). **L)** Quantification for the ratio of colocalization map PSD95/streptavidin objects divided by the total SYP positive objects in the z-stack for cortex and hippocampus images (Huygens colocalization and object analysis, n=4 brains, 6 images per mouse per region). **M)** Quantification of Spearman colocalization coefficient for PSD95/streptavidin in hippocampus and cortex images (Huygens colocalization analysis, n=4 brains, 6 images per mouse per region).

In previous studies from our lab, we reported that BIN1 primarily localizes to presynaptic terminals; however, postsynaptic localization of BIN1 was also observed, albeit to a lesser extent^17^. To determine the degree of biotinylation in either synaptic terminal compartment, we performed immunofluorescence staining of paraffin-embedded sections using the presynaptic marker synaptophysin and the postsynaptic marker PSD95, followed by counterstaining with streptavidin to detect all biotinylated proteins. Layer 1 of the cortex and the *stratum oriens* of the hippocampus were selected for synapse quantification due to high synaptic density and relatively fewer cell bodies. Synaptophysin staining of the cortex revealed abundant puncta as expected, and the streptavidin staining displayed a dense web of biotinylated proteins in the neuropil **(Fig. 3F)**. The overlap between the two is depicted in a colocalization map, showing that 18% of synapses in cortex Layer 1 contained proteins biotinylated by BIN1iso1-TID **(Fig. 3H)**. A similar pattern emerged in the hippocampus, with streptavidin labeling observed in 17.6% of total synapses **(Fig. 3G and 3H).** The Spearman coefficient analysis indicated that the cortex had a coefficient of 0.161, whereas the hippocampus had a coefficient of 0.217 **(Fig. 3I)**. Thus, these two measurements were fairly consistent, with 15-20% of synapses labeled by BIN1iso1- TID. We then stained postsynaptic terminals with PSD95 in layer 1 of the cortex and hippocampus and observed abundant postsynaptic puncta that exhibited moderate overlap with the streptavidin staining of biotinylated proteins in the merged image and the colocalization **(Fig. 3J and 3K)**. Quantification of the overlap revealed that 22.0% and 14.4% of total synapses were biotinylated in the cortex and hippocampus, respectively **(Fig. 3L)**. Moreover, the overlap in labeling of the cortex and hippocampus was quantified to have Spearman coefficients of 0.179 and 0.129, respectively **(Fig. 3M)**, indicating that there are relatively fewer biotinylated postsynaptic terminals in the hippocampus than in the cortex. Interestingly, a comparison of the colocalization maps shows a slightly lower density of biotinylated postsynaptic terminals in the hippocampus compared to presynaptic terminals. Overall, the results from the immunostaining analysis demonstrate that upon expression in mouse brain neurons, BIN1iso1-TID localizes to synapses and promotes the proximity labeling of proteins in both pre- and post-synaptic compartments.

## The mouse brain neuronal BIN1 interactome

We combined the BIN1iso1-TID-mediated proximity biotin-labeled tissue lysates from the cortex and hippocampus, processing the samples as described above **(Fig. 2A)** for label-free quantitative mass spectrometry using DIA. The same data analysis pipeline we used for the N2a dataset was applied for the mouse brain dataset **(Fig. 2B)**. We identified 897 proteins with a z-score >1 as the *in vivo* neuronal BIN1iso1 interactome in the mouse brain. A protein network was generated using the STRING database by inputting a subset of the BIN1 interactome selected based on Z-score >2 enrichment (265 proteins) to visualize the network better **(Fig. 4)**. The edges connecting the nodes were filtered to a high confidence score > 0.7 and the node size corresponds to the -Log of the p-value. The network features several major clusters that are color-coded and annotated according to the most prominent terms among their interconnected nodes. Since the edge threshold is set to a high confidence score, this resulted in 150 singlet nodes and 5 doublets, likely due to the limited information on their functions and functional relationships. Alternatively, this could indicate that this latter group of proteins participates in distinct, non-overlapping cellular functions compared to the rest of the BIN1 interactome. These singlets were color-coded to represent their average BIN1-TID vs. Cyto-TID ratio and were maintained in the network visualization since several of them were top hits from our analysis (RANG and IF2B2, encoded by *Ranbp1* and *Igf2bp2*, respectively).

**Fig 4.**
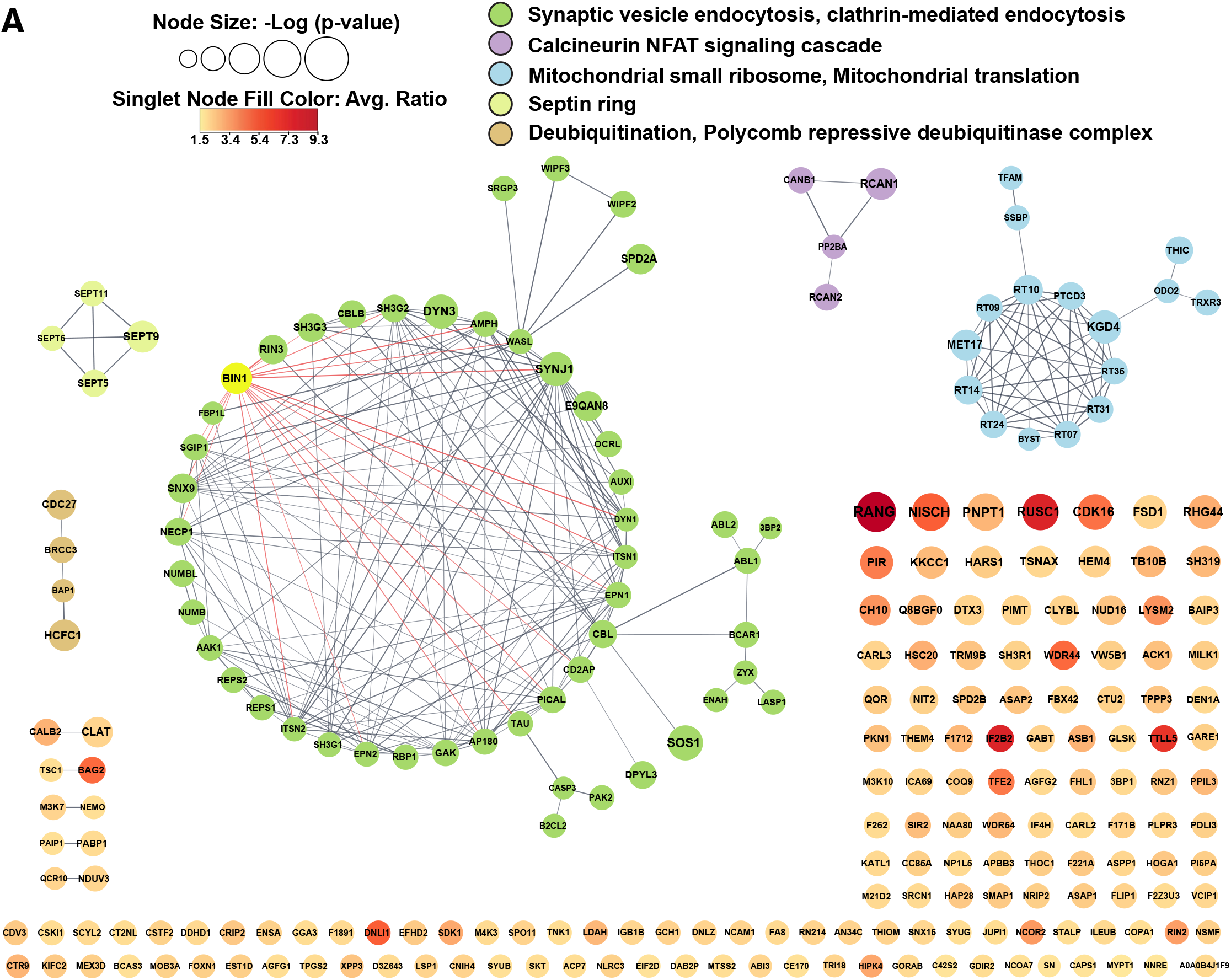
The BIN1iso1-TID interactome in homeostatic mouse brain neurons. **A)** A STRING network was generated for enriched proteins in BIN1iso1-TID mouse brain neurons for z-score >2. High confidence edge threshold score > 0.70. Node size corresponds to the -Log (Welch’s T-test p-value). Nodes are annotated with colors based on the representative terms that represent the group of nodes. Singlet nodes are color-coded with a gradient that corresponds to the average ratio of BIN1-TID vs Cyto-TID.

The most interconnected group of proteins in the BIN1iso1 neuronal interactome is the green cluster (50 nodes, 173 edges), which includes mouse BIN1, indicated in yellow. This cluster is rich in proteins related to synaptic vesicle endocytosis and clathrin-mediated endocytosis. A total of 19 nodes were found directly connected to BIN1 as first neighbors, each located within the green cluster. Many of BIN1’s first neighbors are highly interconnected, as can be seen by the crossover of their edges. One of the top hits in the interactome is the BIN1 homolog AMPH, which, like BIN1, is enriched in nerve terminals and involved in multiple steps in synaptic vesicle recycling^24,71^. Another BIN1 first neighbor is TAU (encoded by *Mapt*, 2.50 average ratio), which stabilizes microtubules in a phosphorylation-dependent manner and localizes to synaptic terminals^72^.

The purple cluster consists of PP2BA and CANB1, the catalytic and regulatory subunits, respectively, of the calcium-dependent calcineurin (CaN) serine/threonine phosphatase that targets a broad range of substrates, including transcription factors (NFAT), microtubule proteins (Tau), and mitochondrial proteins^73^. Interestingly, this cluster also contains RCAN1 and RCAN2 (Regulator of Calcineurin 1 and 2), which can inhibit the catalytic activity of CaN^74^. The yellow cluster features several septin proteins, a group of GTP-binding proteins that form hetero-oligomeric structures serving as scaffolds for the cytoskeleton and are involved in intracellular transport^75^. In rat primary hippocampal neurons, SEPT9 was identified as a dendritic microtubule-associated protein that differentially regulates the motility of axonal and dendritic vesicles through its association with kinesin motors, slowing KIF5 and speeding up KIF1A as they enter dendrites^76^. The small brown cluster contains several proteins involved in ubiquitination pathways, such as BAP1, a deubiquitinating enzyme that regulates nuclear multiprotein complexes responsible for controlling the cell cycle, DNA damage response, and apoptosis, with HCFC1 reported as a substrate^77,78^. Finally, the second largest cluster features 16 nodes and 49 edges corresponding to mitochondrial proteins. This group of proteins primarily relates to the mitochondrial small ribosomal subunit (RT09, RT10, PTCD3, RT35, RT31, RT07, RT24, RT14) and the translation machinery, with 13 out of 16 localizing to the mitochondrial matrix.

A volcano plot depicting the 897 BIN1iso1 interactome proteins with a z-score >1 is shown in **Figure 5A**. Several known BIN1 interacting proteins, such as AMPH1, DYN1, SYNJ1, and ASAP2, are represented in the plot, along with many high-confidence proteins identified in this study as members of the brain *in vivo* BIN1 interactome. The top 30 proteins in the interactome, ranked by their z-score, are depicted in **Figure 5B** with a heat map corresponding to the Log2 transformation of the label-free quantification (LFQ) values for each animal (left) and a horizontal bar graph displaying their average BIN1/Cyto ratio (right). Next to BIN1, AMPH1 (7.82 average ratio) is ranked 4^th^ by the z-score in the BIN1 neuronal interactome and stands out in the heatmap as having high statistical significance **(Fig. 5B)**. Whereas BIN1 can form homodimers^26^, there is evidence suggesting that BIN1 forms heterodimers with AMPH1 and that they can be coimmunoprecipitated from the brain in an equimolar complex^24^. Thus, using an orthogonal approach, BIN1-TID *in vivo* interactome mapping validates previous findings.

**Fig 5.**
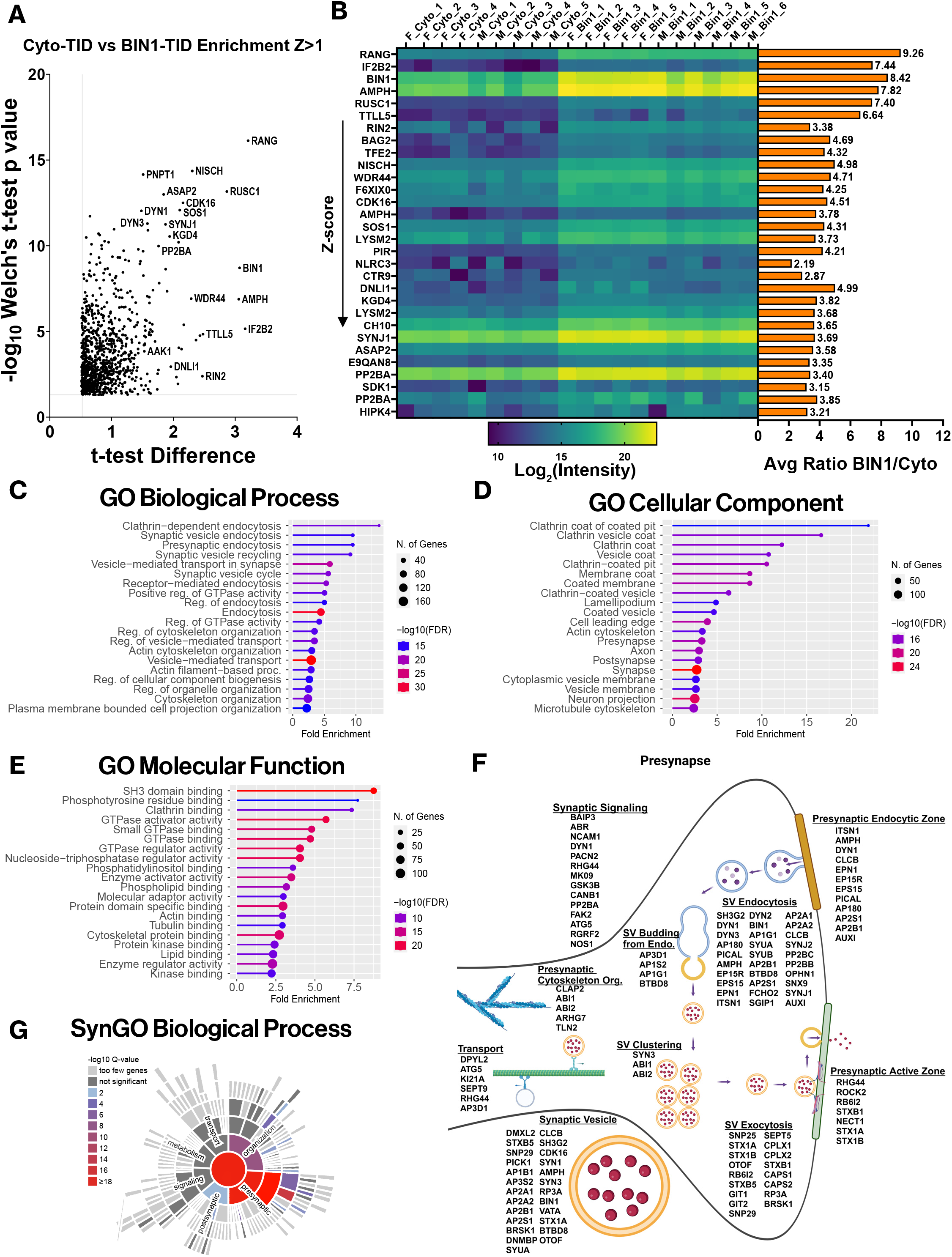
Analysis of the BIN1iso1-TID interactome reveals novel proximal or interacting proteins and functions for BIN1. **A)** Volcano plot displaying T-test Difference vs. - Log10(Welch’s T-test p-value) for the mouse brain neuron BIN1iso1-TID interactome (Z-score >1) **B)** On the left, heat map of Log2(intensity) for top 30 proteins, by descending Z-score and a horizontal bar graph on the right displaying the average ratio BIN1iso1-TID/Cyto-TID. **C)** Lollipop graph of Gene Ontology (GO) term analysis for biological process. Lines and lollipops are colored according to -Log10(FDR) and the sizes of the lollipops correspond to the number of genes with fold enrichment on the x-axis. **D)** GO term analysis for cellular component. Lines and lollipops are colored according to -Log10(FDR) and the sizes of the lollipops correspond to the number of genes with fold enrichment on the x-axis. **E)** GO term analysis for molecular function. Lines and lollipop are colored according to -Log10(FDR) and the sizes of the lollipops correspond to the number of genes with fold enrichment on the x-axis. **F)** Presynaptic SynGO BP and CC terms cartoon for the BIN1iso1-TID interactome. Annotated with abundant terms and proteins from each term detected in interactome z-score >1. **G)** SynGO biological process heat map color coded according to -log10 (Q-value).

Next, we conducted GO analysis of the BIN1iso1 interactome with Z>1 to gain biological insights. The most represented biological pathways included vesicle-mediated transport (Enrichment FDR - 1.67e-33), endocytosis (Enrichment FDR - 6.27e-32), and more specifically vesicle-mediated transport in the synapse (Enrichment FDR - 1.14e-22) **(Fig. 5C)**. As anticipated from the prior characterization of BIN1 localization in synapses^17^, the GO cellular component analysis identified the synapse (Enrichment FDR - 4.2e-27), with the presynapse (Enrichment FDR - 5.58e-18) and postsynapse (Enrichment FDR - 5.30e-16) both highly represented in the dataset **(Fig. 5D)**. The results of GO Molecular Function analysis revealed that SH3 domain binding (Enrichment FDR – 5.56e-24), clathrin binding (Enrichment FDR – 1.52e-9), and cytoskeletal protein binding (Enrichment FDR – 6.83e-18) are prominent in the *in vivo* BIN1 interactome **(Fig. 5E)**. This functional enrichment of BIN1 proximal/interacting proteins are consistent with BIN1’s role in neurotransmitter vesicle dynamics at the synapse and clathrin-mediated endocytosis (primarily defined using non-neuronal cells), highlighting the significance of the cytoskeleton in these processes.

In excitatory neurons, the loss of BIN1 expression hinders neurotransmitter vesicle release rather than vesicle endocytosis^17^. To gain further insights into the specific synaptic functions of proteins identified as the BIN1iso1 interactome, we conducted a Synapse GO (SynGO) analysis with annotations based solely on published experimental evidence curated by experts in synapse biology. In the BIN1iso1 interactome, the Cellular Component analysis reveals that there are 159 synaptic proteins, with 87 located at the presynapse and 86 at the postsynapse (albeit with some overlap between the two synaptic sites) **(Fig. 5F and Supplemental Fig. 2D).** More specifically, the terms for synaptic vesicle (25 of 153 proteins), extrinsic component of presynaptic endocytic zone membrane (9 of 11 proteins; EPN1, EPS15, EPS15R, PICAL, AP180, AMPH, AP2S1, AP2B1, AUXI), and postsynaptic actin cytoskeleton (5 of 21 proteins; FLIP1, SRC8, CTTB2, ABI2, ITSN1) were highly enriched. The SynGO Biological Process analysis supported a role for BIN1 in synaptic vesicle exocytosis, which was significantly enriched (17 of 83 proteins) **(Fig. 5G and 5F)**. BIN1 is in the proximity of multiple steps in synaptic vesicle dynamics as evidenced by BIN1-TID-mediated biotinylation of synaptic vesicle docking (2 of 8 proteins; STXB5, SEPT5), synaptic vesicle priming (8 of 19 proteins; STXB1, RB6I2, CAPS1, CAPS2, RP3A, OTOF, STXB5, BRSK1), and synaptic vesicle fusion to the presynaptic active zone membrane (3 of 6 proteins; STX1B, CPLX1, CPLX2). Thus, the results of the unbiased analysis of the BIN1iso1 interactome in neurons align well with our earlier findings which showed the predominant localization of BIN1 at presynaptic terminals and revealed a non-redundant role for BIN1 in synaptic vesicle release dynamics^17^.

Whereas the first neighbors in the BIN1 interactome we identified have supporting evidence in the literature for their association with BIN1 **(Supplemental Fig. 2B)**, several of the proteins exhibiting the highest average fold ratio in our dataset remain either unlinked from the network as singlets or are separated by several degrees from BIN1 in the network (**Fig. 4**).

Consequently, many proteins in the network have no known physical or functional association with BIN1. One of the most highly enriched proteins is Insulin-like Growth Factor 2 mRNA-binding protein 2 (IF2B2 or formerly IMP2, encoded by *Igf2bp2*; 7.4-fold increase). IF2B2 is an RNA-binding protein characterized as an N^6^-methyladenosine (m^6^A) reader, which stabilizes target RNA, regulates intracellular RNA localization, and regulates their translation in an m^6^A-dependent manner^79^. Knockdown of IF2B2 expression in cultured hippocampal neurons mitigated Alzheimer’s β-amyloid-induced neuronal damage and pyroptosis, partly by downregulating m^6^A-mediated NLRP expression and inflammasome formation^80^. Inhibiting IF2B2 in acute myeloid leukemia demonstrated anti-leukemic effects both *in vitro* and *in vivo* while also regulating expression of MYC^81^, a well-documented interacting partner of BIN1, which, in this context, functions as a tumor suppressor ^82–85^. However, like many other proteins in the BIN1iso1 neuronal interactome, this study is the first to connect IF2B2 to BIN1, and the functional implications of the putative interaction have yet to be determined.

### Concordance analysis of the *in vitro* and *in vivo* BIN1iso1 interactomes

As described above, we have applied the TurboID labeling strategy to identify several novel and overlapping BIN1 interacting and proximal proteins both *in vitro* in the simple N2a culture system and *in vivo* in mouse brain neurons. We conducted a combined analysis of the two interactome datasets to explore the concordant findings and nominate the high-confidence BIN1 interactome proteins for validation. We identified 92 proteins shared between the two datasets (**Fig. 6A)** and created a STRING network to determine their functional relationships with this subset. Since these proteins were replicated across the two datasets, we adjusted the edge score threshold to a medium confidence score of > 0.4 to better predict potential PPI relationships. To effectively visualize the quantitative contribution of individual membership, we color-coded the network nodes according to their fold ratio in the N2a BIN1iso1 interactome and displayed the size of the nodes based on their fold ratio in the BIN1iso1 mouse brain interactome **(Fig. 6B)**. In the network, over half of the BIN1 interactome is grouped into 3 major clusters. Cluster 1, the largest cluster, contains BIN1 and is enriched in proteins that function in neurotransmitter vesicle dynamics (SH3G1, FCHO2, BTBD8, SYNJ1, SYNJ2), vesicle-mediated transport (SNX18, SH3G1, SYNRG, FCHO2, and WIPF2), regulation of GTPase activity (ASAP1, ASAP2, and SNX18), and protein ubiquitination (CBL, CBLB, CDC27, and ITCH). Within this cluster, BIN1 is directly connected to only 9 of 24 the proteins based on information available in the STRING and DAVID databases **(Fig. 6B, left)**. The second-largest cluster, Cluster 3, is enriched in mitochondrial proteins that predominantly localize to the mitochondrial matrix and are involved in mitochondrial translation (RF1M, RT31, RT35, RT09, RT27, PTCD3, RT10, RT24, RT14) **(Fig. 6B, right)**. These proteins are prt of the mitochondrial small ribosomal subunit complex. In addition, this cluster included a group of proteins that function in the citric acid cycle and respiratory electron transport (NDUV2, NDUV3, NDUF7, ODP2, ODO2, and ODO1). This cluster is particularly notable since these proteins were reproducibly labeled by proximity biotinylation under both *in vitro* and *in vivo* conditions, yet, BIN1 has no known mitochondrial association. Cluster 2 comprises nucleoplasmic proteins linked to histone modification (BAP1, HCFC1, SET1A, YETS2, KANL3) and, more specifically, the H4 histone acetyltransferase complex (HCFC1, YETS2, KANL3) **(Fig. 6B, middle)**. BIN1 does not have a defined role in histone modification. In addition to the three clusters, 29 singlet proteins and 4 doublets were left unconnected to any other nodes in the network based on the information available in the literature, despite being proximity biotinylated by BIN1iso1-TID *in vitro* and *in vivo*.

**Fig. 6.**
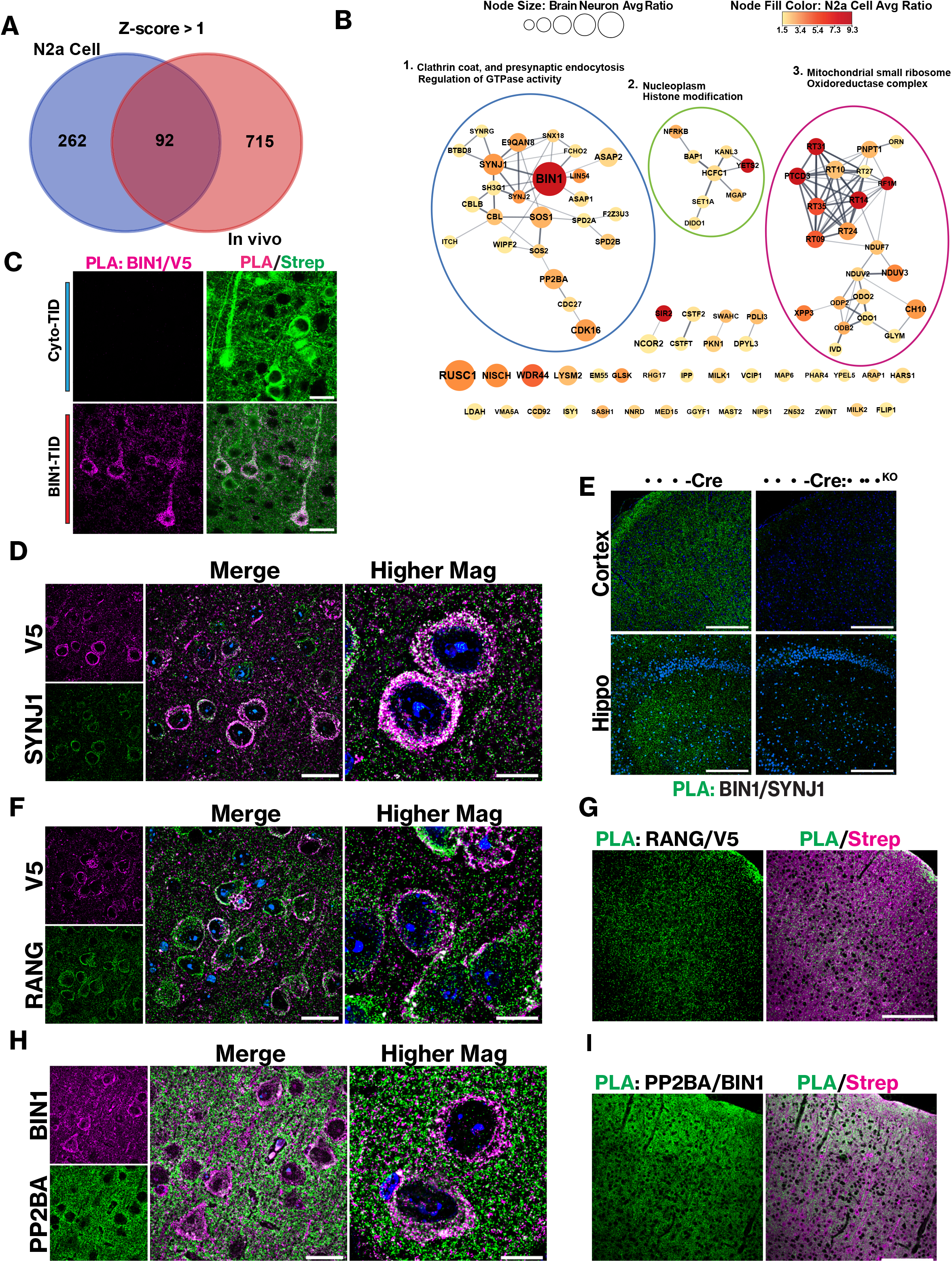
Validation of BIN1iso1 N2a cell and mouse brain neuron interactome top hits. **A)** Venn diagram showing the N2a cell BIN1 interactome overlap with the BIN1 mouse brain neuron interactome (z-score >1). **B)** A STRING network was generated for enriched proteins in 92 common proteins between N2a BIN1iso1-TID and BIN1iso1-TID mouse brain neurons for z- score >1. A medium confidence edge threshold score > 0.40. Node size corresponds to the BIN1iso1-TID mouse brain interactome. Singlet nodes are color-coded with a gradient that corresponds to the average ratio of the N2a BIN1-TID interactome. Nodes are annotated with terms and selections based on the representative terms that highlight the group of nodes. **C)** Proximity ligation assay (PLA) of Cyto-TID vs BIN1iso1-TID using anti-V5 and anti-BIN1. Puncta represent proteins within a 40 nm molecular distance. Co-stained with streptavidin to display biotinylated proteins (scale bar = 25 µm). **D)** IF of BIN1iso1-TID brain using anti-SYNJ1 and anti-V5. Overlap of images (scale bar = 25 µm). Higher magnification images are z-stacks projected as a Sum (scale bar = 10 µm). **E)** PLA using anti-BIN1 and SYNJ1 in brains of neuron-specific Emx-Cre:*Bin1* knockout and Emx-Cre control mice. The Emx-Cre:*Bin1* knockout mouse lacks BIN1 expression in neurons and oligodendrocytes in the forebrain, thus serving as a negative control for PLA. **F)** IF of BIN1iso1-TID brain using anti-RANG and anti-V5. Overlap of images (scale bar = 25 µm). Higher magnification images are z-stacks projected as a Sum (scale bar = 10 µm). **G)** PLA of BIN1iso1-TID using anti-V5 and anti-RANG. Co-stained with streptavidin to display biotinylated proteins **H)** IF of BIN1iso1-TID brain using anti-PP2BA and anti-BIN1. Overlap of images (scale bar = 25 µm). Higher magnification images are z-stacks projected as a Sum (scale bar = 10 µm). **I)** PLA of BIN1iso1-TID using anti-PP2BA and anti- BIN1. Co-stained with streptavidin to display biotinylated proteins for reference.

## Proximity ligation assay (PLA) validation select BIN1iso1 interactome proteins

We conducted a series of PLAs to validate select proteins in the BIN1 interactome. First, we carried out a control experiment where PLA was performed on brain sections from mice transduced with rAAV expressing BIN1iso1-TID or Cyto-TID, using antibodies against V5 and BIN1, followed by streptavidin staining. We reasoned that since both the BIN1 and V5 antibodies would label BIN1isoTID, a positive PLA signal would be generated, which would overlap with the streptavidin staining. In contrast, little, if any, PLA signal is anticipated in mice expressing Cyto-TID, as endogenous BIN1 (labeled only by the BIN1 antibody) is unlikely to be within a molecular distance of cytosolic Cyto-TID (labeled by the V5 antibody). As expected, we readily observed PLA signals in the neuronal cell bodies, apical dendrites, and neuropil in the cortex and hippocampus of animals transduced with rAAV expressing BIN1iso1-TID **(Fig. 6C and Supplemental Fig. 2C)**. In contrast, PLA signals were largely absent in the brains of animals transduced with rAAV expressing Cyto-TID, as predicted.

Next, we decided to validate select highly enriched proteins in the *in vivo* dataset or both *in vivo* and *in vitro* datasets. As a positive control, we chose a known BIN1 interacting protein, SYNJ1, which was biotinylated by BIN1iso1-TID and enriched in both datasets (7.5-fold in N2a cells and 4.25-fold in mouse neurons) **(Fig. 6B)**. SYNJ1 is a phosphatase with two phosphoinositide phosphatase domains and a proline-rich domain, is known to bind to BIN1 and other proteins with BAR and SH3 domains such as AMPH1 and endophilin^86^. Notably, BIN1 has also been known to interact with AMPH1 and endophilin^61,87^, and our neuronal BIN1 interactome includes AMPH1 and members of the endophilin A family (SH3G1, SH3G2, and SH3G3) as enriched interactors **(Fig.4)**. SYNJ1 plays established roles in endocytosis, clathrin uncoating during clathrin-mediated endocytosis, and the regulation of phosphatidylinositol^88^.

Consequently, there has been an intense focus on SYNJ1 and its specific functions in synaptic vesicle recycling and trafficking. Co-immunostaining analysis of SYNJ1 and BIN1iso1-TID revealed a substantial overlap of SYNJ1 and BIN1 in the soma, in addition to numerous puncta in the neuropil of the cortex and hippocampus **(Fig. 6D and Supplemental Fig. 3A)**. We wanted to confirm that endogenous mouse BIN1 is proximal to SYNJ1 in the mouse brain. To this end, we performed PLA using antibodies against SYNJ1 and BIN1; the latter recognized a shared epitope present in all BIN1 isoforms and is known to immunostain BIN1 in neurons and glia^40^. To demonstrate the specificity of BIN1 and SYNJ1 PLA labeling under the conditions employed in our study, we conducted PLA in neuron-specific *Emx*-Cre:BIN1 knockout and *Emx*- Cre control mice. The *Emx*-Cre:*Bin1* knockout mouse lacks BIN1 expression in excitatory neurons and oligodendrocytes in the forebrain^17^, thus serving as a negative control for BIN1- SYNJ1 PLA. The results indicate that endogenous mouse BIN1 is close to SYNJ1, as evidenced by the robust PLA signal, and that the absence of BIN1 in excitatory neurons significantly diminishes the PLA signal in the cortex and hippocampus of *Emx*-Cre:*Bin1* knockout brains **(Fig. 6E)**. These results validate SYNJ1 as a *bona fide* BIN1 interacting protein in neurons.

Next, we focused on RANG (*RanBP1)*, as it was the most highly enriched protein in the mouse brain BIN1iso1 interactome (9.26-fold over Cyto-TID; z-score 6.58) and a singlet within the BIN1 interactome network **(Figs. 4 and 5B)**. RANG is primarily known for its role in nucleocytoplasmic transport, by regulating RAGP1-mediated hydrolysis of RanGTP to RanGDP, which releases cargo for export from the nucleus via exportins or allows cargo binding by importins for protein import into the nucleus^89,90^. RANG and BIN1iso1-TID were observed to co- localize in the neuronal cell body, dendrites, and neuropil of the cortex in mice transduced with rAAV expressing BIN1iso1-TID **(Fig. 6F),** as well as in the hippocampus **(Supplementary** Fig. 3C). PLA using anti-V5 and anti-RANG antibodies exhibited abundant positive puncta throughout the cortex, localized to the periphery of the neuronal cell body and the neuropil, as seen in the overlay with streptavidin staining, which reveals all proximity biotinylated proteins **(Fig. 6G)**. In the hippocampus, positive puncta are present in the pyramidal layer and in the *stratum oriens* and *stratum radiatum* **(Supplementary** Fig. 3D).

Additionally, we selected PP2BA, the catalytic subunit of calcineurin phosphatase, which was identified in both BIN1 interactome datasets (5.87-fold in N2a cells and 3.40-fold in mouse neurons). Calcineurin depends on calcium to dephosphorylate its target proteins^91^. In the presynaptic terminal, calcineurin is involved in both endocytosis and exocytosis. Blocking calcium influx, calmodulin, or the pharmacological inhibition of calcineurin blocks endocytosis, while the extent of calcium influx regulates the speed of both rapid and slow endocytosis by calcineurin dephosphorylating endocytic proteins such as AMPH1, SYNJ1, and DYN1^92,93^. In rAAV-transduced mice expressing BIN1iso1-TID, PP2BA co-localized with BIN1iso1-TID as well as endogenous BIN1, as visualized by immunostaining with an anti-BIN1 antibody, in the neuronal cell body and overlapping puncta in the neuropil **(Fig. 6H)**. In the hippocampus, the PP2BA signal is particularly pronounced in the *stratum radiatum*, where many synapses are located **(Supplemental Fig. 3E)**. The BIN1 signal overlaps with PP2BA in the neuronal soma, dendrites, and neuropil. PLA showed robust positive labeling in the neuronal soma and synapses, especially abundant in areas of the cortex rich in synapses, such as layer 1 **(Fig. 6I)**. In agreement, the PLA signal in the synapse-rich *stratum oriens* and *stratum radiatum* of the hippocampus is notably substantial **(Supplementary** Fig. 3F). The congruent findings from *in vitro* and *in vivo* proximity labeling, along with PLA validation of SYNJ1, RANG, and PP2BA underscore the reliability of the BIN1 interactome identified in this study.

## Phosphorylation site analysis reveals AAK1 as a key BIN1 interacting partner that facilitates endocytosis at the synapse

Kinases function as molecular switches in many biological processes, and BIN1 itself undergoes phosphorylation^24,39^. We aimed to characterize the phosphorylation landscape in the BIN1iso1-TID-labeled proteins to gain additional insights into BIN1 function and the kinases that might phosphorylate BIN1. We conducted a phosphorylation site search using existing mass spectrometry data from streptavidin-enriched BIN1iso1-TID mouse samples, which were not prepared for conventional phosphoproteomics workflow. Nevertheless, we identified 252 phosphorylated proteins in the BIN1iso1-TID proteomics dataset. To refine our focus, we filtered the phosphorylated proteins based on the mouse brain BIN1iso1-TID dataset, selecting only those with a significant z-score >1, resulting in 72 proteins. We then constructed a STRING network using proteins corresponding to a z-score >2, with their corresponding phospho-sites annotated to the nodes **(Fig. 7A)**. While we did not detect phosphorylated BIN1 in this dataset, we did find phosphor-sites on BIN1 homolog AMPH1 (1 p-site), SYNJ1 (2 p-sites), DYN3 (1 p-site), AP-2-associated protein kinase 1, AAK1 (4 p-sites), and TAU (7 p-sites). Many of these proteins have biological functions linked to vesicle-mediated transport, endocytosis, and synaptic vesicle dynamics, and they localize to pre- and post-synaptic sites, axons, and vesicles.

**Fig 7.**
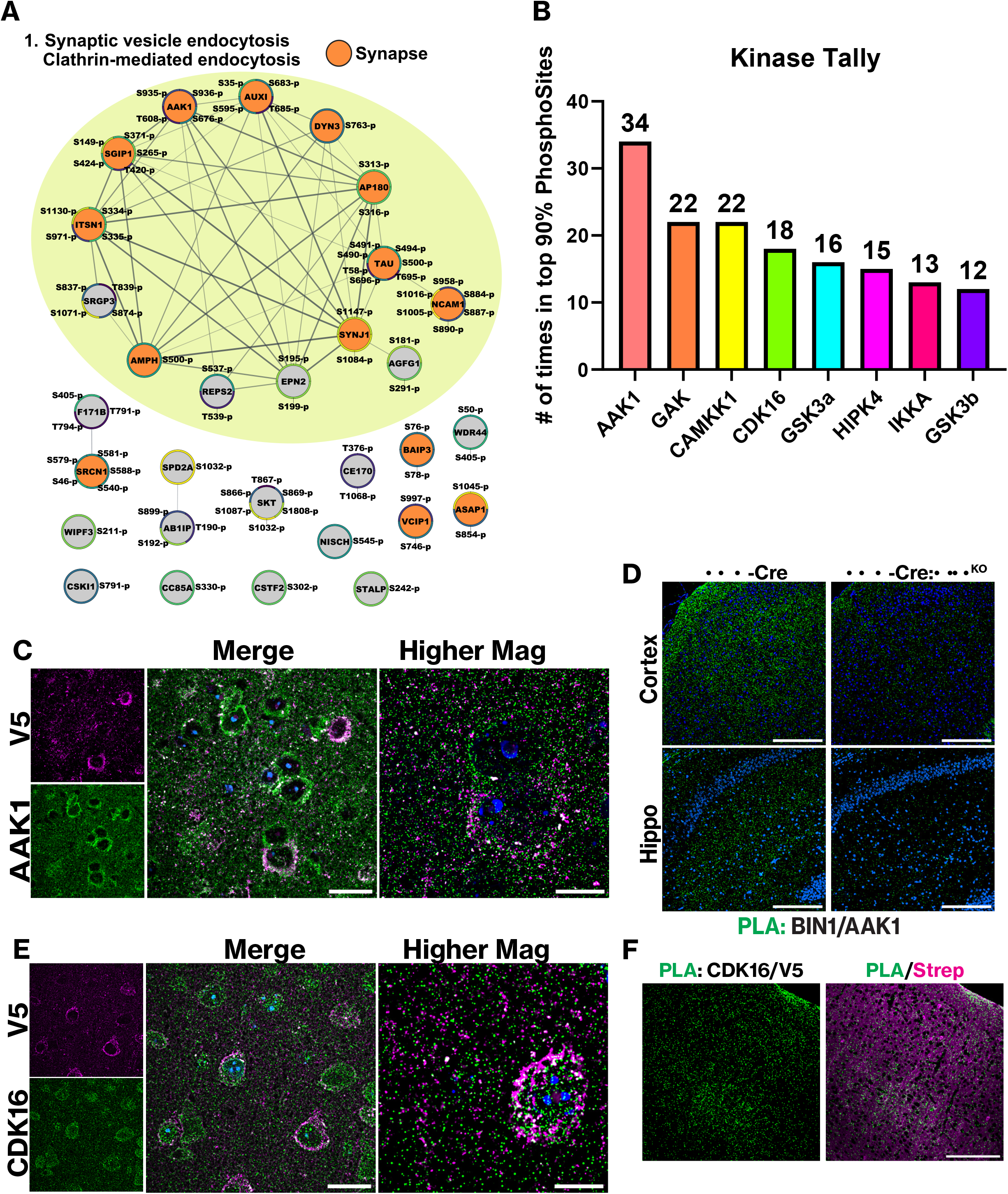
Phosphorylation site analysis of BIN1iso1-TID neuronal interactome identifies AAK1 and CDK16. **A)** A STRING network was generated for proteins identified through the phosphorylation site analysis of BIN1iso1-TID mouse brain neurons for z-score >2. A medium confidence edge threshold score > 0.40 was used. Protein nodes have the detected phosphorylation sites mapped to them. Proteins that localize to the synapse are colored in orange while the annotated group with green background corresponds to endocytic proteins. **B)** Using PhosophoSitePlus prediction. This displays the probability for a specific kinase to phosphorylate the target protein at each experimentally observed phosphorylated site in the interactome. We set >90% as the threshold for inclusion of a particular kinase in our results. The kinases were tallied for phosphoproteins filtered from the BIN1iso1-TID interactome (Z-score >2). **C)** IF of BIN1iso1-TID brain using anti-AAK1 and anti-V5. Overlap of low mag images (scale bar = 25 µm). Higher magnification images are z-stacks projected as a Sum (scale bar = 10 µm). **D)** PLA using anti-AAK1 and anti-BIN1 antibodies. Mice are neuron-specific Emx- Cre:BIN1 knockout and Emx-Cre control mice. The Emx-Cre:*Bin1* knockout mouse lacks BIN1 expression in neurons and oligodendrocytes in the forebrain, thus serving as a negative control for PLA. **E)** IF of BIN1iso1-TID brain using anti-CDK16 and anti-V5 antibodies. Overlap of low mag images (scale bar = 25 µm). Higher magnification images are z-stacks projected as a Sum (scale bar = 10 µm). **F)** PLA of BIN1iso1-TID using anti-CDK16 and anti-V5 antibodies. Co- stained with streptavidin to display biotinylated proteins for reference.

To start understanding which kinase in the BIN1 interactome might be important for BIN1 functions, we used PhosphoSitePlus and its kinase prediction database to predict which kinase may be responsible for each phosphosite in the high-confidence phosphoproteins with a z-score >2 in the neuronal BIN1 interactome **(Fig. 7A)**. Since phosphorylation at a specific site within a protein can potentially be mediated by multiple kinases, the PhosphoSitePlus kinase prediction shows the probability of a given kinase phosphorylating the target protein at each experimentally observed phosphorylated site in our interactome. We set >90% as the threshold for including a particular kinase in our results and found that AAK1 was predicted to phosphorylate BIN1 interacting proteins at 34 sites **(Fig. 7B)**. In addition, CDK16, identified in both BIN1iso1 *in vitro* and *in vivo* datasets, was predicted in this analysis to potentially phosphorylate the BIN1 interactome at 18 sites. Therefore, we decided to validate AAK1 and CDK16 as BIN1 proximal proteins.

AAK1 is a serine/threonine kinase originally identified for its phosphorylation of the µ2 subunit of the AP-2 complex, and it is highly enriched at presynaptic terminals in neurons^94^. In peripheral cells, it colocalizes with clathrin and AP-2 and is found at the leading edge of migrating cells. Additionally, phosphorylation of the µ2 subunit of AP-2 decreases transferrin internalization^94^. Indeed, overexpression of AAK1 impairs transferrin endocytosis, whereas knockdown of AAK1 expression impairs transferrin recycling from the early endosome, suggesting that AAK1 is crucial for multiple steps in the endosomal pathway^95^. Through immunofluorescence labeling, we observed that AAK1 and BIN1iso1-TID colocalize in punctate structures within the neuronal soma and neuropil in the cortex and hippocampus **(Fig. 7C and Supplemental Fig. 3G)**. Further proximity analysis via PLA using AAK1 and BIN1 antibodies showed strong PLA signals in the mouse brain cortex; we observed a marked decrease in the PLA signal in the *Emx*-Cre:*Bin1* knockout brain **(Fig. 7D)**. These findings demonstrate the proximity of endogenous BIN1 to AAK1. Considering the proteomic identification of AAK1 as the most abundant kinase in synaptic vesicles^96^, and its shared role with BIN1 in regulating synaptic vesicle dynamics^17,96^ our identification of AAK1 as a BIN1 proximity protein located within molecular distance raises the possibility of a functional interaction between them at the presynaptic terminal.

Cyclin-dependent kinase 16 (CDK16, previously known as PICTAIRE1) was highly enriched in both BIN1iso1 datasets (7.1-fold in N2a and 4.5-fold in the mouse brain interactome). CDK16 is a Ser/Thr kinase involved in spermatogenesis, cell cycle progression, neurite outgrowth, and vesicle trafficking^97^. Currently, there is no known connection between BIN1 and CDK16.

Through immunostaining, we observed considerable overlap between BIN1iso1-TID and CDK16 in the pyramidal neuron cell body and the neuropil of the cortex and hippocampus **(Fig. 7E and Supplementary** Fig. 3I). PLA using V5 and CDK16 antibodies confirmed their localization at a molecular distance within the neuronal soma and in the neuropil **(Fig. 7F)**. The PLA puncta in the hippocampus were primarily found in the *stratum radiatum* and *stratum oriens*, regions with a high density of synapses **(Supplementary** Fig. 3J). These findings validate AAK1 and CDK16 as potential BIN1 interacting kinases that may regulate neuronal BIN1 function directly and/or through phosphorylation of BIN1 interactome proteins involved in the endocytic pathway.

## Discussion

### BIN1’s proximity to synaptic endocytosis and exocytosis

As a member of the BAR domain family of proteins, BIN1 was initially believed to have a redundant role in endocytosis. The identification of *BIN1* as the second most common risk allele for LOAD has sparked an interest in exploring its function in neurons and other cell types in the brain. Previous studies, primarily based on affinity pull-down assays and candidate approaches, identified fewer than 35 BIN-interacting proteins^9^. Our unbiased proteomics strategy, facilitated by proximity biotin labeling, has significantly broadened the BIN1 interactome by uncovering 361 proteins in N2a cells and a much larger group of 897 proteins in the hippocampal and cortical neurons of the mouse brain. A greater number of BIN1 interactome proteins were found at neuronal synapses, consistent with BIN1 localization at the synapse. Intriguingly, the BIN1 interactome also encompassed proteins localized to the mitochondria, potentially indicating BIN1’s function in cellular metabolism beyond membrane remodeling and cytoskeletal associations.

In a previous study, we reported that the loss of BIN1 in excitatory neurons leads to deficits in vesicular release probability, characterized by an increased number of docked vesicles and a buildup of the reserve pool of synaptic vesicles^17^. These findings suggested that BIN1 function promotes efficient neurotransmitter vesicle release and dynamics at the presynaptic sites in hippocampal excitatory neurons undergoing sustained neuronal activity. The SynGO analysis of the *in vivo* BIN1iso1 neuronal interactome supports BIN1’s role in synaptic vesicle release and predicts its involvement in endocytosis. Notably, the BIN1 interactome comprised 159 synaptic proteins, divided between the presynaptic terminal (87 proteins) and postsynaptic terminal (86 proteins), many which reside at the pre- and post-synaptic endocytic zone and in synaptic vesicles. In terms of biological function, the BIN1 interactome was particularly enriched for the synaptic vesicle cycle and for functions related to synaptic vesicle endocytosis, including proteins that regulate synaptic vesicle budding from endosomes (4 of 4 proteins in this function; AP3D1, AP1S2, AP1G1, and BTBD8), which is also a clathrin-dependent process. In addition, synaptic vesicle exocytosis was significantly enriched (17 of 83 proteins involved in this pathway). There are several defined steps in the synaptic vesicle exocytosis pathway during which vesicles from the reserve pool are loaded with neurotransmitters and sequentially are docked, primed, and then undergo fusion to release their cargo^98^. BIN1 is proximal during each of these processes: synaptic vesicle docking (2 of 8 proteins; STXBP5, and SEPTIN5), synaptic vesicle priming (8 of 19 proteins; STXBP1, ERC1, CADPS, CADPS2, RPH3A, OTOF, STXBP5, and BRSK1), and synaptic vesicle fusion with the presynaptic active zone membrane (3 of 6 proteins; STX1B, CPLX1, and CPLX2). The SynGO analysis indicates that BIN1 plays a crucial role in synaptic vesicle recycling at both the endocytic and exocytic interfaces, supporting and expanding upon our previous findings from the knockout study^17^. While the BIN1iso1 interactome in neurons and the SynGO analysis define BIN1’s protein neighborhood, this does not necessarily imply BIN1 interacts directly with each protein. Consequently, the interactome dataset we present offers numerous opportunities for hypothesis-driven follow-up studies to further clarify BIN1’s mechanistic role at the synapse.

## Phosphorylation site analysis uncovers potential regulatory networks for BIN1 interactors

Although BIN1’s involvement in membrane remodeling and vesicular dynamics is increasingly recognized, relatively little is known about how BIN1 dynamically regulates these processes. We reasoned that dynamic phosphorylation could influence BIN1 function and conducted a phosphorylation site analysis of the BIN1 interactome proteins. Then, we examined the phosphosites within each high-confidence interactome member, to predict kinases that might be responsible for their phosphorylation. The most notable kinase from this analysis was AAK1, which is linked to endocytosis through the phosphorylation of the µ2 subunit of the AP-2 complex^94,99^. A recent study investigated the temporal dynamics of clathrin-coated vesicle formation resulting from AP-2 phosphorylation and found that phosphorylation of µ2 at T156 by AAK1 switches the AP-2 complex to an open conformation, facilitating interaction with clathrin, NECAP (part of the BIN1 interactome), and BIN1 as the clathrin-coated pit matures, allowing BIN1 to recruit dynamin for clathrin-coated vesicle scission^99^. Through PLA, we confirmed that AAK1 localizes within molecular distance to endogenous BIN1 in both the neuronal soma and the neuropil. While we did not identify phosphorylated peptides derived from BIN1 in our analysis, BIN1 is known to be phosphorylated at several sites, including 9 within its clathrin and AP-2 (CLAP)-binding domain. Therefore, it would be interesting to investigate whether AAK1 phosphorylates BIN1 or influences the AP-2:BIN1 interaction. CDK16 is another kinase significantly enriched in both the N2a and mouse neuron BIN1 interactome, validated by PLA. CDK16 is involved in various cellular processes such as the cell cycle, vesicle trafficking, spindle orientation, skeletal myogenesis, neurite outgrowth, secretory cargo transport, and cell growth and proliferation, among others^97^. An *in vitro* chemical genetic screen using mouse brain lysates identified AAK1, dynamin 1, and SYNJ1 as substrates of CDK16^100^. Thus, CDK16 kinase represents another potential regulator of the physiological functions of BIN1 interactome proteins throughout multiple stages of the endocytic pathway.

## BIN1’s potential roles in the cell cycle, histone modification, and mitochondria

We were surprised that some of the most enriched proteins in the BIN1 interactome datasets weren’t related to endocytosis or vesicle trafficking. In the BIN1 N2a cell interactome, none of the top 10 hits, with the exception of mouse endogenous BIN1, were involved in endocytosis. Proteins associated with the cell cycle, histone modification, and mitochondrial translation machinery were among the most enriched in the N2a dataset, and many of these proteins were also found in the mouse brain neuron dataset.

Our data suggests that a hitherto unknown BIN1 function may mediate the import and export of proteins into and out of the nucleus, accounting for the large number of nuclear proteins associated with the cell cycle and histone modification represented in the BIN1 interactome. RANG was the most highly enriched protein from the mouse brain neuronal BIN1 interactome. Only a few studies have focused on RANG in mature neurons, with most showing significant roles for RANG in neuron development, including migration of neural crest cells^101^, axon specification, dendritic arborization, and cortical neuron polarity^102^. In peripheral sensory neurons exposed to axonal injury, RANG was locally translated at the injury site, where it facilitated the hydrolysis of RanGTP and enabled the formation of importin signaling complexes (importin-beta, importin alpha, and dynein) for retrograde injury signaling^103^. BIN1 has established roles in vesicle-mediated transport, and our study found that it is close to RNA-binding proteins such as IF2B2 involved in localized translation^104^. RANG, RANB3, and IMA4 were significantly enriched in the mouse neuron BIN1 interactome, and RAN was enriched in the N2a BIN1 interactome. These findings raise the possibility that BIN1 may play a role in the Ran-associated import of cargo proteins to the nucleus and in retrograde signaling from synaptic terminals to respond to stimuli at distant synapses.

Another surprising finding was the number of mitochondrial proteins in the BIN1 interactome. A majority of them are part of the mitochondrial ribosome, specifically the small mitochondrial ribosomal subunit. A small subset of proteins contributes to mitochondrial functions related to cellular respiration and the TCA cycle. Like most mitochondrial proteins, the mitoribosomal subunit genes are transcribed in the nucleus, translated in the cytosol, and then imported into the mitochondria, where they assemble into the small ribosomal subunit^105^. Many of the mitochondrial proteins in our interactome dataset localize to the mitochondrial matrix.

There are several possibilities for these observations: a) BIN1 may localize to the mitochondrial membrane or mitochondria-associated membranes where it labels imported proteins; b) BIN1 might be near cytosolic ribosomes that are translating mitochondrial proteins, labeling them as the *de novo* proteins are imported into the mitochondria; and c) since BIN1 plays an important role in the scission of endosomes, it may assists DYN2 in mitochondrial fission. BIN1 has not been described as having a role in mitochondrial biology or localized to mitochondria. Since BIN1 hasn’t been localized to mitochondria, we first checked whether any BAR domain superfamily member associates with mitochondria. One BAR protein, FAM92A1, was found to localize to mitochondrial membranes and required for mitochondrial ultrastructure and function^106^. Nevertheless, our efforts with high-resolution imaging failed to demonstrate mitochondrial localization of BIN1 in cultured N2a cells and the brain. Another reason BIN1 might be in close proximity to mitochondria is its association with dynamin 2, which is involved in mitochondrial fission^107^. Due to BIN1’s interaction with dynamin proteins, there is a possibility that BIN1 will be recruited during this process. Additionally, one protein of interest is TFAM, a nuclear-encoded mitochondrial transcription factor that recognizes promoter sites on mtDNA and recruits other initiation factors^108^. TFAM contains an SH3 binding motif, suggesting a potential direct interaction with BIN1. If BIN1 is a possible regulator of mtDNA transcription, aberrant BIN1 expression in diseases such as AD could lead to mitochondrial dysfunction.

Overall, the diversity of the BIN1 interactome indicates that BIN1 may be involved in multiple processes, including gene transcription, nuclear protein transport, and mitochondrial biology, with wide-ranging impacts on cell health and biology.

## Insights on Alzheimer’s disease

As the second most common risk factor for LOAD, extensive efforts have been made to determine how BIN1 influences LOAD risk. The BIN1 interactome described in this study may provide insights into how BIN1 and its protein partners may influence the disease. We compared the GWAS LOAD risk gene proteins to the BIN1iso1 interactome and found that BIN1 was proximal to 16 LOAD risk proteins: CRADD, ICA69, BCAS3, TAU, CD2AP, PKHA1, ABI3, CCDC6, NCK2, TNIP1, PICAL, GAB2, RIN3, FAK2, ANR55, and TLN2. Most of these proteins regulate the cytoskeleton, endocytosis, and transport. Several of these proteins have been linked to BIN1 in AD amyloid or Tau pathology investigations. Two recent *in vitro* studies suggested that BIN1 plays a role in Aβ generation, one through BIN1 interaction with BACE1^109^ and the other through BIN1 interaction with RIN3^35,110^. Our proximity labeling study supports an interaction between BIN1 and RIN3, whereas BACE1 was not identified in the BIN1 interactome. Although *in vitro* evidence has shown that BIN1’s SH3 domain can bind to Tau^36–38^, the interaction is weakened by Tau phosphorylation^37^. Thus, it is not clear whether an interaction between BIN1 and Tau provides a mechanistic explanation for findings from studies implicating BIN1 in the propagation of Tau pathology^33,40,111–115^. An equally plausible explanation is that BIN1’s role in neurotransmitter vesicle release^17^ and synaptic vesicle dynamics influences the efficiency of Tau spreading. TAU becomes hyperphosphorylated in the brains of individuals with AD, causing it to aggregate and lose its ability to stabilize microtubules and leading to neuronal dysfunction^116^. Phosphorylation site analysis of our proteomics data identified 7 phosphorylated sites on Tau (T58p, S490p, S491p, S494p, S500p, T695p, and S696p). The identification of GSK3A, GSK3B, calcineurin, and SIR2 in the BIN1 interactome raises the possibility that as an adaptor protein, BIN1 might functionally interact with these regulators of Tau phosphorylation.

## Conclusions

We employed TurboID proximity biotinylation coupled with proteomics to elucidate the neuronal interactome of BIN1, a significant susceptibility risk factor for LOAD. Our analysis provided unbiased *in vivo* validation of previously reported BIN1-interacting proteins and revealed hundreds of previously unknown proximal or interacting proteins, laying the foundation for future investigations into the neuronal functions of BIN1. Elucidating the BIN1 interactome confirmed BIN1’s previously established roles in synaptic vesicle dynamics, endocytosis, and cytoskeletal association, while also uncovering additional potential roles for BIN1 in mitochondrial translation, metabolism, and nuclear histone modification. We validated that AAK1, CDK16, SYNJ1, PP2BA, and RANG are *bona fide* BIN1 proximal proteins in brain neurons that may regulate BIN1 or other critical proteins within BIN1’s regulatory network. Overall, our data positions BIN1 at the nexus of many essential cellular processes and suggests novel avenues for investigating how BIN1 elevates LOAD risk.

## Data Availability

The mass spectrometry data have been deposited to the ProteomeXchange Consortium via the PRIDE^117^ partner repository with the dataset identifier PXD060638.

## Supplemental Data

This article contains supplemental data.

## Acknowledgements

We would like to thank Om Patel for his help in optimizing antibody dilutions for use in paraffin-embedded mouse brains.

## Author Contributions

Conceptualization, J.D.M., S.M.S., G.T.; Methodology, J.D.M., S.W., J.W., J.G., S.M.S.; Investigation, J.D.M., J.W., J.G.; Formal Analysis, J.D.M., J.G., S.M.S.; Visualization, J.D.M., S.M.S., G.T.; Writing – Original Draft, J.D.M., J.G., S.M.S., G.T.; Writing – Review & Editing, J.D.M., S.W., J.W., J.G., S.M.S., G.T.; Supervision, S.M.S., G.T.; Project Administration, G.T.; Funding Acquisition, J.D.M., S.M.S., G.T.

## Abbreviations

Bridging Integrator 1 (BIN1), Late-Onset Alzheimer’s Disease (LOAD), Alzheimer’s disease (AD), BIN1/amphiphysin/RVS167 (BAR), Clathrin-AP-binding (CLAP), MYC-binding domain (MBD), SRC homology 3 (SH3), TurboID (TID), proline-rich domains (PRD), neuro-2a cells (N2a), label-free quantification (LFQ), data-independent acquisition (DIA), false-discovery rate (FDR), protein-protein interaction (PPI), Gene Ontology (GO), recombinant adeno-associated virus (rAAV), proximity ligation assays (PLA), immunofluorescence (IF), Search Tool for the Retrieval of Interacting Genes/Proteins (STRING)

---

**Supplemental Figure 1.**
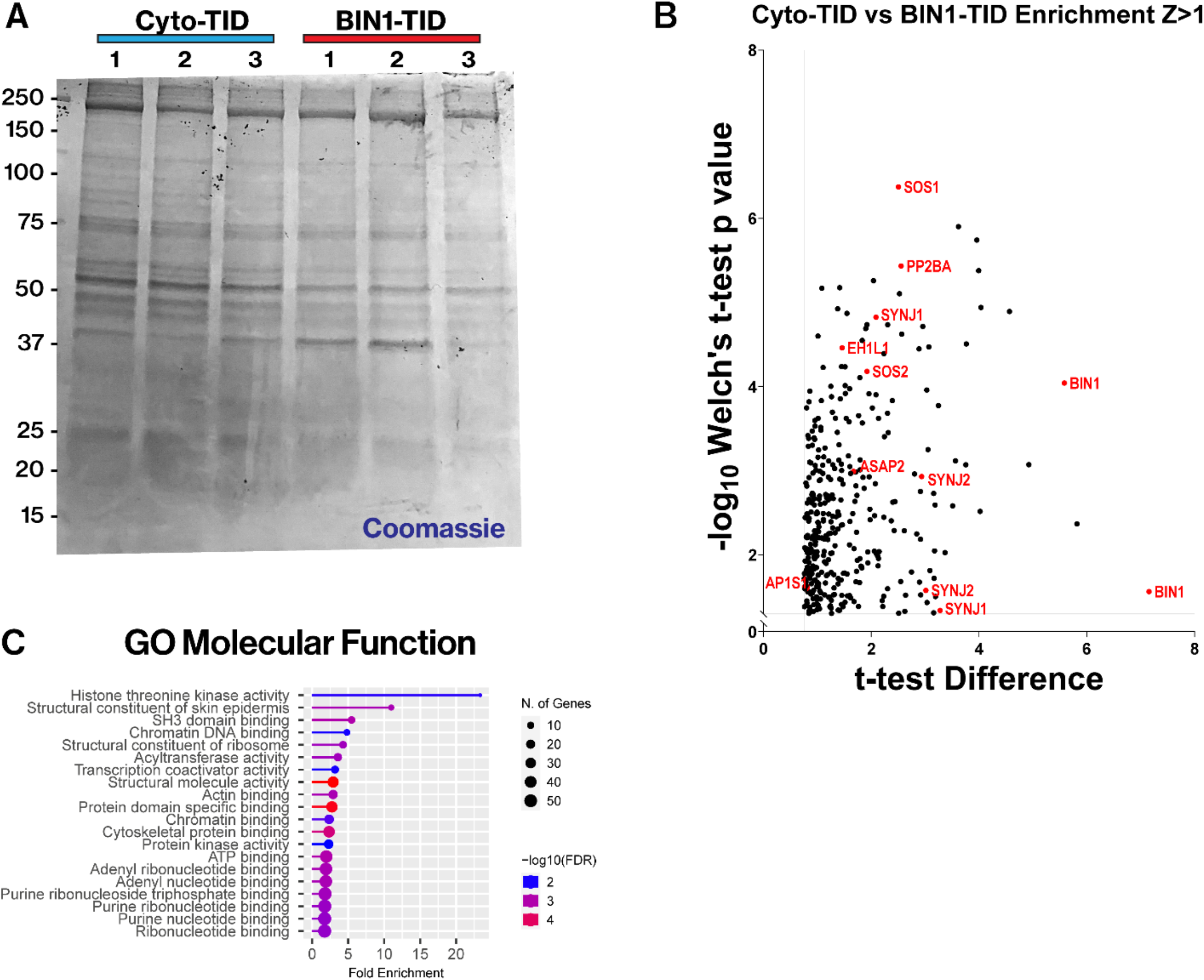
N**2**a **cells stably expressing BIN1-TID label known BIN1- associated proteins and molecular functions. A)** Streptavidin magnetic beads captured biotinylated protein from stable N2a cells stably expressing Cyto-TID and BIN1-TID. Protein eluted from the beads, run on a gel and total protein stained with Coomassie. **B)** Volcano plot displaying the N2a BIN1-TID positive protein hits for z-score >1 and previously reported BIN1- associated proteins labeled in red, t-test difference (x>0.758) vs. -log_10_ Welch’s t-test p-value (y>1.301). **C)** GO molecular function (FDR <0.05), node size (number of genes), node color (- log_10_FDR) and fold enrichment on the x-axis.

**Supplemental Figure 2.**
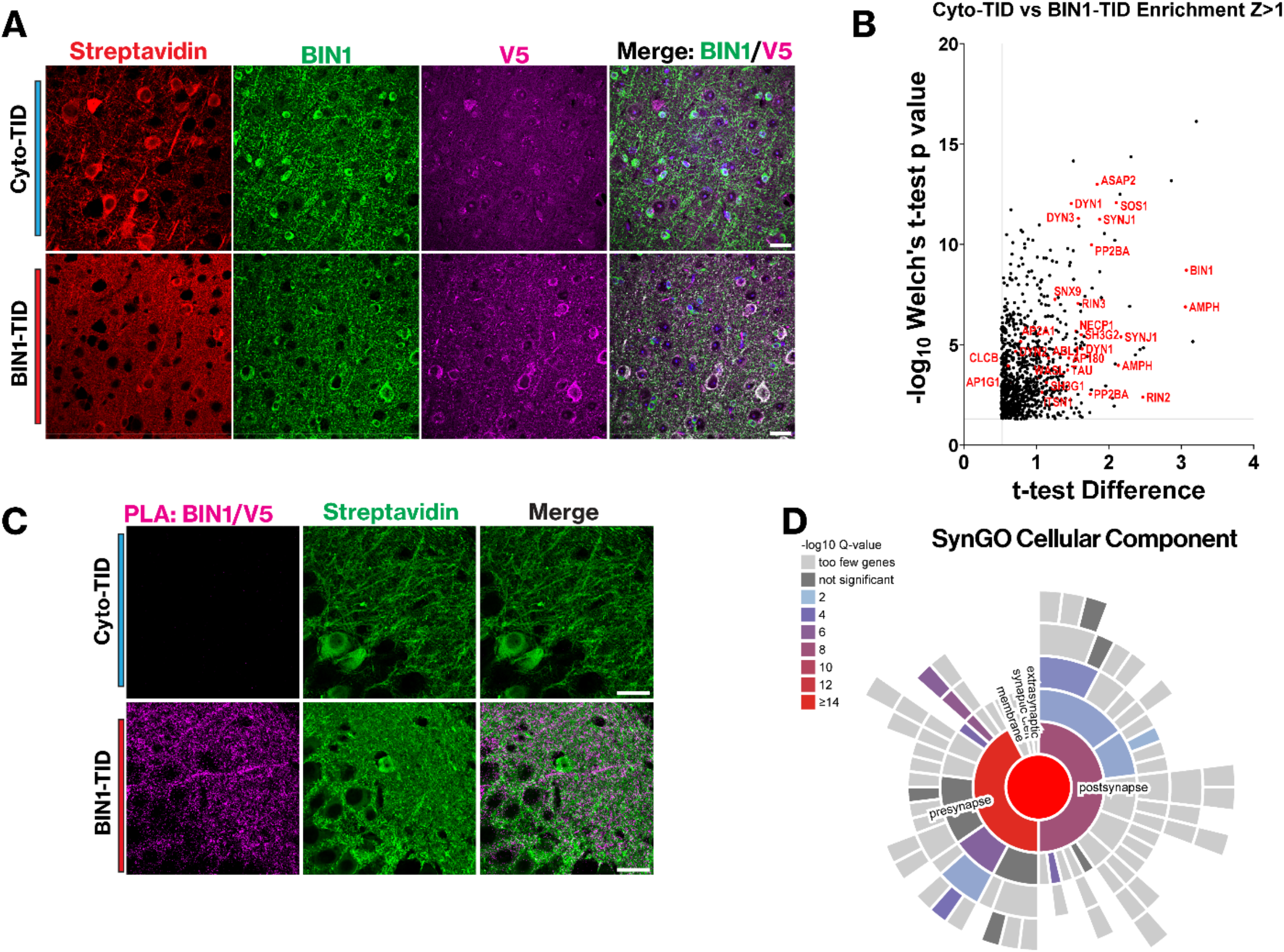
B**I**N1**-TID colocalizes with endogenous mouse BIN1 and biotinylates presynaptic proteins A)** IF staining of mouse cortex using Anti-BIN1, anti-V5, and streptavidin. The merge showing the overlap of V5 and BIN1 (scale bar = 25µm) **B)** Volcano plot displaying the mouse brain neuron BIN1-TID positive protein hits for z-score >1 and previously reported BIN1-associated proteins labeled in red, t-test difference (x>0.526) vs. -log_10_ Welch’s t-test p-value (y>1.301). **C)** PLA using primary antibodies against V5 and BIN1. Counterstained with streptavidinin green (scale bar = 25 µm) **D)** SynGO cellullar component analysis (Q-value < 0.05) color correspondning to -log_10_Q-value and annotated with synaptic compartments showing preferential enrichment of the BIN1-TID interactome in the presynaptic terminal.

**Supplemental Figure 3.**
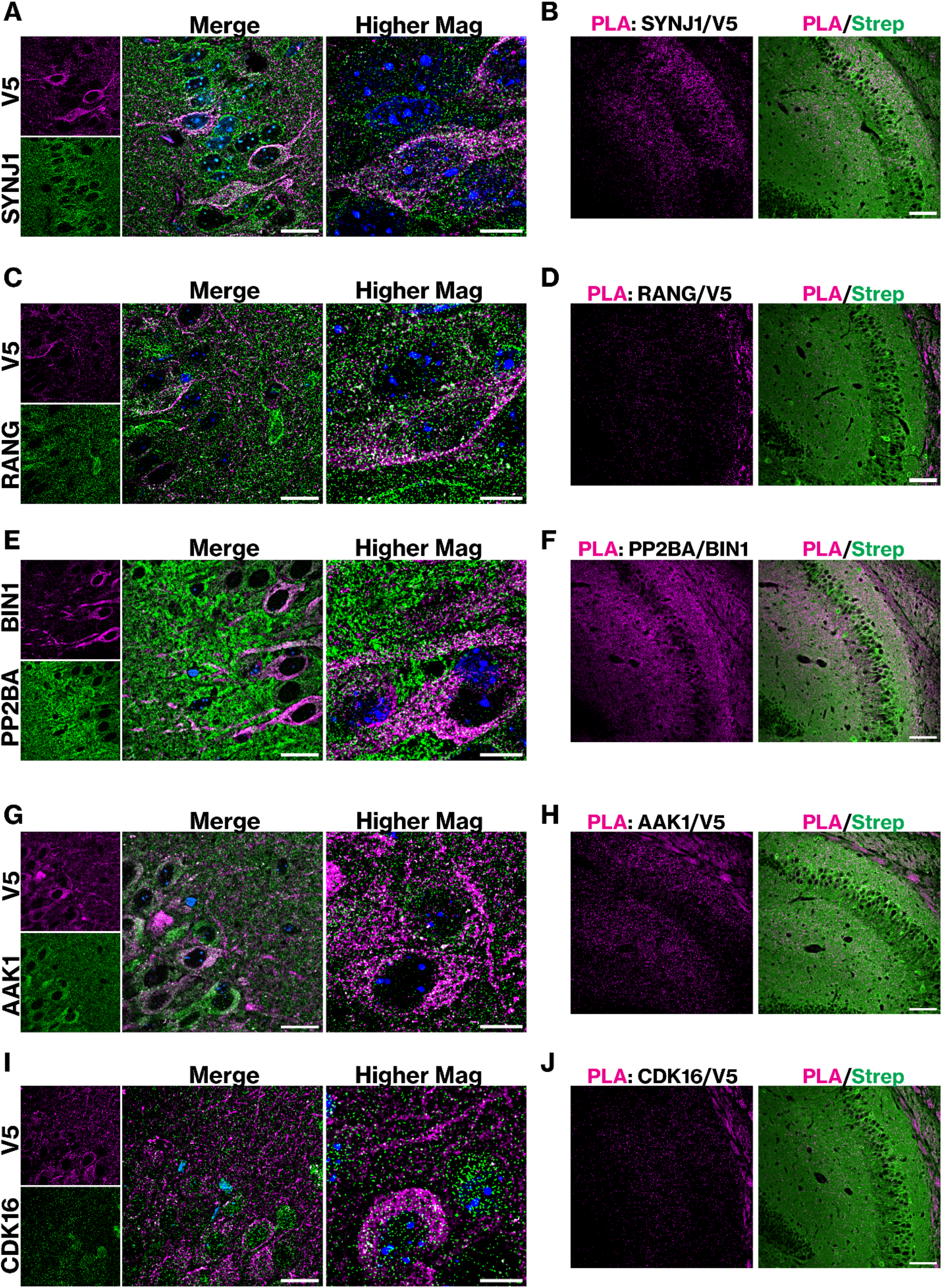
B**I**N1**-TID is within a molecular distance of SYNJ1, RANG, PP2BA, AAK1, and CDK16 in the hippocampus. A)** IF of BIN1iso1-TID hippocampus using anti- SYNJ1 and anti-V5. Overlap of images (scale bar = 25 µm). Higher magnification images are z-stacks projected as a Sum (scale bar = 10 µm) **B)** PLA using anti-V5 and -SYNJ1. The merge shows PLA signal and streptavidin staining of biotinylated protein for spatial reference (scale bar = 200 µm). **C)** IF of BIN1iso1-TID hippocampus using anti-RANG and anti-V5. Overlap of images (scale bar = 25 µm). Higher magnification images are z-stacks projected as a Sum (scale bar = 10 µm). **D)** PLA of BIN1iso1-TID hippocampus using anti-V5 and anti-RANG. Co-stained with streptavidin to display biotinylated proteins (scale bar = 200 µm) **E)** IF of BIN1iso1- TID hippocampus using anti-PP2BA and anti-BIN1. Overlap of images (scale bar = 25 µm). Higher magnification images are z-stacks projected as a Sum (scale bar = 10 µm). **F)** PLA of BIN1iso1-TID using anti-PP2BA and anti-BIN1. Co-stained with streptavidin to display biotinylated proteins for reference (scale bar = 200 µm). **G)** IF of BIN1iso1-TID hippocampus using anti-AAK1 and anti-V5. Overlap of low mag images (scale bar = 25 µm). Higher magnification images are z-stacks projected as a Sum (scale bar = 10 µm). **H)** PLA using anti- AAK1 and anti-V5 antibodies. Co-stained with streptavidin to display biotinylated proteins for reference (scale bar = 200 µm). **I)** IF of BIN1iso1-TID hippocampus using anti-CDK16 and anti- V5 antibodies. Overlap of low mag images (scale bar = 25 µm). Higher magnification images are z-stacks projected as a Sum (scale bar = 10 µm). **J)** PLA of BIN1iso1-TID using anti-CDK16 and anti-V5 antibodies. Co-stained with streptavidin to display biotinylated proteins for reference (scale bar = 200 µm).

